# A new type of ERGIC-ERES membrane contact mediated by TMED9 and SEC12 is required for autophagosome biogenesis

**DOI:** 10.1101/2021.05.19.444771

**Authors:** Shulin Li, Rui Yan, Jialu Xu, Shiqun Zhao, Xinyu Ma, Qiming Sun, Min Zhang, Ying Li, Jun-Jie Gogo Liu, Liangyi Chen, Sai Li, Ke Xu, Liang Ge

**Affiliations:** State Key Laboratory of Membrane Biology, Beijing, China; Tsinghua-Peking Center for Life Sciences, Beijing, 100084, China; School of Life Sciences, Tsinghua University, Beijing, 100084, China; Department of Chemistry, University of California, Berkeley, CA, USA; Beijing Advanced Innovation Center for Structural Biology, Beijing, 100084, China; Beijing Key Laboratory of Cardiometabolic Molecular Medicine, Institute of Molecular Medicine, Peking University, Beijing, 100871, China; National Center for Nanoscience and Technology, Beijing, 100871, China; Department of Biochemistry, Department of Cardiology of Second Affiliated Hospital, Zhejiang University School of Medicine, Hangzhou, 310058, China

**Keywords:** Autophagy, autophagosome, ERGIC, ERES, TMED9, SEC12, LC3, COPII, membrane contact, ER, Golgi, phagophore, PAS, TFG, FIP200

## Abstract

Under stress, the endomembrane system undergoes reorganization to support autophagosome biogenesis, which is a central step in autophagy. How the endomembrane system remodels has been poorly understood. Here we identify a new type of membrane contact formed between the ER-Golgi intermediate compartment (ERGIC) and the ER-exit site (ERES) in the ER-Golgi system, which is essential for promoting autophagosome biogenesis induced by different stress stimuli. The ERGIC-ERES contact is established by the interaction between TMED9 and SEC12 which generates a short distance opposition (as close as 2-5 nm) between the two compartments. The tight membrane contact allows the ERES-located SEC12 to transactivate COPII assembly on the ERGIC. In addition, a portion of SEC12 also relocates to the ERGIC. Through both mechanisms, the ERGIC-ERES contact promotes formation of the ERGIC-derived COPII vesicle, a membrane precursor of the autophagosome. The ERGIC-ERES contact is physically and functionally different from the TFG-mediated ERGIC-ERES adjunction involved in secretory protein transport, and therefore defines a unique endomembrane structure generated upon stress conditions for autophagic membrane formation.

## INTRODUCTION

Autophagy is a bulk cytoplasmic degradation process, which is mediated by the lysosome and is involved in numerous physiological processes and pathological conditions^1,2^. The regulation of autophagy is achieved by a cascade signal relay of autophagy-related (ATG) proteins. First, the serine/threonine protein kinase complex, consisting of ULK1/2, FIP200, ATG13, and ATG101, forms a protein scaffold, likely through phase transition, at the phagophore assembly site (PAS)^1–4^. Second, the class III phosphatidylinositol (PI)-3 kinase complex I, composed of ATG14L, Beclin-1, VPS34, and P150, is recruited to the ER-associated membranes and catalyzes the formation of PI3-phosphate (PI3P)^1,2,4^. Third, the PI3P effector WIPI2, together with FIP200, recruits ATG16L and the protein conjugate ATG5-ATG12, which acts as an E3 in a ubiquitin-like conjugation process and, together with ATG7 (E1) and ATG3 (E2), catalyzes the covalent linkage of LC3 proteins to phosphatidylethanolamine (PE)^5–11^. The membranes with lipidated LC3s coalesce with ATG9-positive vesicles to build the cup-shaped phagophore, which expands through acquisition of lipids from multiple sources via membrane fusion, ATG2-mediated lipid transfer, and lipid synthesis, in a cradle localized in the ER (omegasome)^12–23^. The phagophore engulfs cytoplasmic contents and closes via ESCRT-mediated membrane scission to form a double-membrane autophagosome^24,25^. The autophagosome finally fuses with a lysosome, or with an endosome and a lysosome, to form an autolysosome and complete the degradation of the enclosed components^1,4^. After degradation, new lysosomes are generated through lysosome-reformation, characterized by tubulation of the autolysosome ^26^.

A central step in autophagy is the formation of the double-membrane autophagosome. The process requires membrane and lipid contribution from the endomembrane system^12^. Although efforts have been made to uncover the membrane sources of the autophagosome in past decades, a definitive answer about which endomembrane compartment generates the autophagic membrane is still lacking, especially concerning the question of what designates the compartment as a membrane source for the autophagosome^4^. Recently, we and others identified the ER-Golgi intermediate compartment (ERGIC) as a key membrane source for autophagosome biogenesis in response to different autophagy stimuli^15,27–32^. However, which molecular components of the ERGIC determine its contribution to autophagosome biogenesis was unknown.

Membrane contact sites are defined as areas of close apposition (usually 10-30 nm in distance) formed between intracellular membrane compartments, in which membrane fusion is absent^33–35^. Multiple types of membrane contact have been identified. For example, the ER has been shown to form diverse membrane contacts with different organelles including mitochondria, the plasma membrane, and peroxisomes^33–36^. These different types of membrane contacts have been shown to regulate inter-organelle communication and dynamics via controlling calcium signaling, lipid transfer, and membrane remodeling^33–35^. However, a holistic view regarding the regulation and function of membrane contact is still lacking, pending the revelation of new types of membrane contacts, what functions the membrane contact sites perform, and how these functions are carried out^33^.

Recent evidence indicates that membrane contacts regulate autophagosome biogenesis. The membrane contact sites of ER-mitochondria and ER-plasma membrane were shown as cradles for PI3P synthesis and autophagosome assembly, likely acting as a PAS^37,38^. On these ER-related membrane contact sites, a dynamic membrane association between the growing phagophore and the ER was indicated to regulate growth, maturation, and transport of the autophagosome^39,40^. Although a link between membrane contact for autophagosomal membrane assembly and maturation has been established, it is unclear if and how membrane contacts regulate the bona fide generation of autophagosomal membrane precursors from the endomembrane system, which has been poorly characterized compared to the process of phagophore assembly.

Here, using cell-free lipidation, immuno-isolation, and mass spectrometry, we identified the ERGIC protein TMED9 as a key regulator of autophagosome biogenesis under multiple stimuli in both non-selective and selective autophagy. The ERGIC-localized TMED9 associates with the ERES protein SEC12, which establishes a dynamic and short distance (as close as 2-5 nm) membrane contact between the ERGIC and the ERES. The ERGIC-ERES contact regulates the generation of ERGIC-derived COPII vesicles (ERGIC-COPII), a new type of COPII vesicle that we have previously shown to be specific for autophagosome biogenesis^29^. The ERGIC-ERES contact is physically and functionally distinct from the TFG-mediated ERES-ERGIC association^41–43^. Therefore, we have identified a new type of membrane contact in the ER-Golgi system regulating the biogenesis of membrane precursors of the autophagosome.

## RESULTS

### Identification of TMED9 as an ERGIC determinant for autophagosome biogenesis

Previously, we developed a cell-free LC3 lipidation assay which allowed us to identify the ERGIC as a most active compartment triggering LC3 lipidation^27^. However, it has been unclear which components of the ERGIC account for its high lipidation activity. Three transmembrane proteins, ATG9, VMP1, and TMEM41B, were shown to be involved in autophagy, in which VMP1 and TMED41B functionally compensate for each other^44–48^. Nonetheless, depletion of either ATG9 or VMP1 in the membrane by different siRNAs did not affect LC3 lipidation in the cell-free assay (Fig. S1A, B). This is consistent with studies showing that depletion of three proteins did not affect lipidation in the cell^45–49^. Actually, three recent studies demonstrated that the three transmembrane proteins act as scramblases that coordinate lipid acquisition in phagophore assembly and growth^50–52^. Therefore, these three proteins are unlikely to contribute to the high LC3 lipidation activity on the ERGIC in autophagy. To look for the unknown active components of the ERGIC, we performed trypsin digestion in combination with Na_2_CO_3_ treatment of the ERGIC and determined LC3 lipidation (Fig. 1A). Na_2_CO_3_ treatment alone, which disrupts membrane structure and washes off peripheral membrane proteins did not affect the LC3 lipidation on the ERGIC (Fig. 1A). Interestingly, trypsin digestion dose-dependently eliminated LC3 lipidation (Fig. 1A). The data indicate that membrane-anchored proteins on the ERGIC account for the high LC3 lipidation activity.

**Figure 1.**
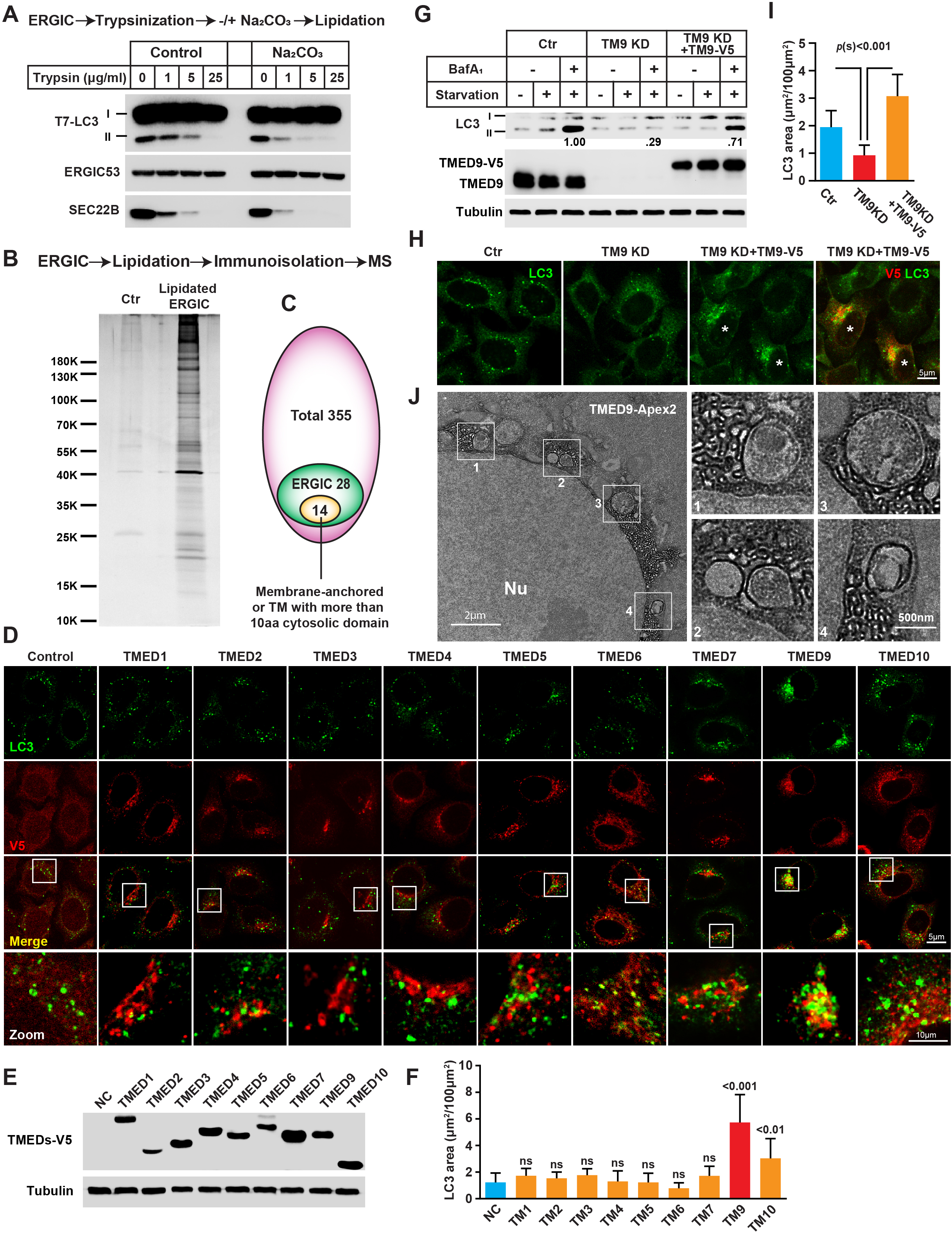
The ERGIC membrane protein TMED9 regulates autophagosome biogenesis. (A) The ERGIC fraction from Atg5KO MEF cells was collected and digested with the indicated concentrations of trypsin with or without Na_2_CO_3_ (0.25 M, pH 11) on ice for 20 min. PMSF (1 mM) was incubated to quench trypsin digestion, and the digested membrane was harvested by 100,000×*g* centrifugation. Cell-free lipidation was performed with the digested membranes and cytosols prepared from starved HEK293T cells. (B) The LC3-lipidated ERGIC was immunoisolated and mass spectrometry was employed to identify proteins enriched in lipidated ERGIC. (C) Venn diagram showing membrane-anchored proteins enriched in the ERGIC. (D) Immunofluorescence of HeLa cells stably expressing TMEDs-V5 after Earle’s Balanced Salf Solution (EBSS) starvation (1h) with V5 and LC3 antibody (E) Immunoblots showing expression of the TMEDs indicated in (D). (F) Quantification of the LC3 puncta area in (D). Error bars represent standard deviations of > 150 cells from three independent experiments (> 50 cells per experiment). P-values were obtained from two-tailed t-test. (G) LC3 lipidation in HeLa cells transfected with control or siRNAs against TMED9 with or without TMED9-V5 re-expression. The cells were incubated in nutrient-rich medium or starved in EBSS in the absence or presence of 500 nM bafilomycin A1 for 1 h. Immunoblots were performed to determine the levels of indicated proteins. Quantification was based on the ratio of lipidated LC3 to tubulin with the control set as 1.00 (control siRNA with starvation and bafilomycin A1). The blots are representative of at least three independent experiments. (H) Immunofluorescence of HeLa cells (control, TMED9 knockdown (KD) or TMED9 KD with TMED9-V5 expression) with anti-V5 and LC3 antibodies. The cells were starved in EBSS for 1 h. Asterisks indicate cells with TMED9-V5 expression. (I) Quantification of the LC3 puncta area (μm^2^/100μm^2^ cell area) analyzed in (H). Error bars represent standard deviations of > 150 cells from three independent experiments (> 50 cell per experiment). P-value was obtained from two-tailed t-test. (J) EM images showing the TMED9-Apex2 labeled membrane and the adjacent autophagosomes. The cells were starved in EBSS for 1 h. Scale bar sizes are indicated in the picture.

We hypothesized that the proteins should be ERGIC-resident membrane proteins and should be able to associate with cytosolic components. Therefore, the cytosolic part of the membrane protein should be long enough to accommodate this association (we set a standard of at least 10 aa in the cytosolic domain). To identify the ERGIC membrane proteins, we performed LC3 lipidation with ERGIC and immunoisolated LC3-labeled ERGIC membrane followed by mass spectrometry analysis. Of the 355 proteins identified, 28 were ERGIC-enriched proteins, of which 14 were transmembrane proteins with cytosolic domains longer than 10 aa (Fig. 1B, C and Table S1). Of the 14 candidates, seven were ERGIC-resident transmembrane emp24 domain-containing (TMED) proteins^53^ (Table S1). To determine the role of TMED proteins in autophagosome biogenesis, we established cell lines separately expressing each of the 9 TMED family members found in humans. Of the 9 TMED proteins, expression of TMED9 increased endogenous LC3 puncta formation (> 3-fold), which is characteristic of the autophagosome (Fig. 1D-F). In addition, knockdown of TMED9, as opposed to any of the other indicated TMED, compromised LC3 lipidation in the cell-free assay (Fig. S1C), indicating that TMED9 solely accounts for LC3 lipidation activity, and is likely involved in autophagosome biogenesis.

To confirm the involvement of TMED9 in regulating autophagosome biogenesis, we performed RNAi-mediated TMED9 knockdown followed by re-expression of TMED9. Depletion of TMED9 abolished starvation-induced LC3 lipidation (∼3.4-fold decrease with starvation and bafilomycin A1), LC3 puncta formation (∼2-fold decrease with starvation), and autophagic vacuoles (∼4-fold decrease with starvation) determined by immunoblot, immunofluorescence, and electron microscopy (EM) respectively (Fig. 1G-I; Fig. S1D, E). CRISPR/CAS9-mediated TMED9 knockout in HeLa cells showed consistent dependence of autophagosome biogenesis on TMED9 as determined by LC3 lipidation (Fig. S1F, G). TMED9 RNAi selectively affected LC3 puncta formation instead of the PAS as revealed by FIP200 puncta indicating that TMED9 is specifically involved in the LC3 lipidation step (Fig. S1H, I). The deficiency of autophagosome biogenesis was reversed by restoring TMED9 expression (Fig. 1G-I; Fig. S1G). We noticed that in TMED9-expressed cells, LC3 puncta distribution was affected and the puncta tended to localize around the TMED9-positive perinuclear compartments (Fig. 1H). The association of autophagosomes with the TMED9 compartment was confirmed by EM analysis, in which double-membrane autophagosomes tightly associated with the TMED9-Apex2-labeled tubulovesicular membrane, a morphology characteristic of the ERGIC (Fig. 1J). The data together indicate that TMED9 is required for autophagosome biogenesis and the TMED9-positive ERGIC compartment may directly contribute membranes to the autophagosome.

### TMED9 regulates autophagosome biogenesis and selective autophagy in response to different stimuli

Autophagy is controlled by different signaling regulators including mTORC1, AMPK, and growth factors^54,55^. We employed different ways of inducing autophagy, including starvation in Earle’s Balanced Salt Solution (EBSS), low glucose, serum removal, and rapamycin treatment, to recapitulate autophagy activation via different signaling regulators (Fig. 2A-F). As shown in Figure 2, depletion of TMED9 compromises autophagosome biogenesis (as shown by LC3 lipidation, Fig. 2A-C) and autophagic flux (as shown by the tandem fluorescence LC3 assay^56^, Fig. 2D-F) induced by different stimuli. Therefore, TMED9-regulated autophagosome biogenesis is a convergent step downstream of different autophagy-regulating signal cascades.

**Figure 2.**
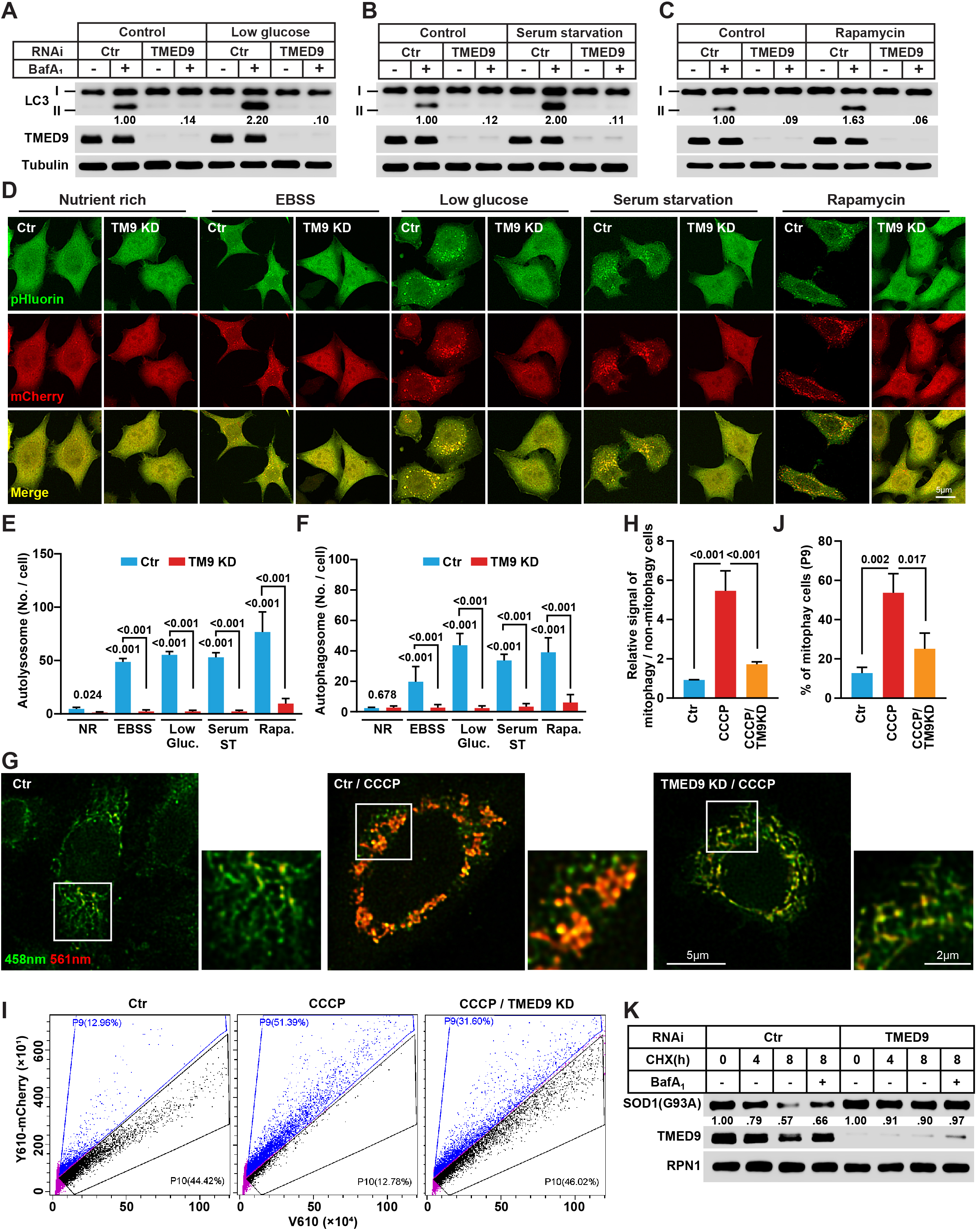
TMED9 is required for multiple types of autophagy in response to different stimuli. (A-C) LC3 lipidation in HeLa cells transfected with control or siRNAs against TMED9. The cells were incubated in nutrient-rich medium, low glucose medium (A), medium without serum (B), or treated with 1 μM rapamycin for 8 h (C) in the absence or presence of 500 nM bafilomycin A1 for 1 h. Immunoblots were performed to determine the levels of indicated proteins. Quantification was performed similarly to Fig.1G. The blots are representative of at least three independent experiments. (D) HeLa cells stably expressing the mCherry-pHluorin-LC3B were transfected with control or siRNAs against TMED9 and treated were incubated in nutrient-rich medium, EBSS (1 h), low glucose medium (8 h), medium without serum (8 h), or treated with 1 μM Rapamycin for 8 h. Confocal microscopy was performed. (E, F) Quantification of autophagosome (yellow, (E)) and autolysosome (red, (F)) numbers per cell as shown in (D). Error bars represent standard deviations of >300 cells from three independent experiments (>100 cell per experiment). P-value was obtained from two-tailed t-test. (G) Control and TMED9 KD HeLa cells were co-transfected with mito-Keima and Parkin and treated with or without 20 µM CCCP for 4 h. Confocal microscopy was performed. The results are representative of at least three independent experiments. (H) The ratio of 568 nm (mitophagy)/440 nm (non-mitophagy) signals was calculated in (G) as relative mt-Keima signal. Error bars represent standard deviations of >300 cells from three independent experiments (>100 cell per experiment). P-value was obtained from two-tailed t-test. (I) Control and TMED9 KD HeLa cells co-expressing mt-Keima and Parkin were treated with or without 20 µM CCCP for 4 h. Cells were analyzed by FACS using V610 and Y610-mCherry detectors (Beckman CytoFLEX LX). The FACS results are representative of at least three independent experiments. (J) The percentage of cells with mitophagy (P9) based on Y610-mCherry/V610 calculated in (I). Error bars represent standard deviations of 3 experiments. P-value was obtained from two-tailed t-test. (K) Lysosome-dependent turnover of SOD1(G93A) in CHX chase assay in control or TMED9 KD HeLa. Quantification was based on the ratio of SOD1(G93A) to RPN1 with the control set as 1.00 (control siRNA at time 0). The blots are representative of at least three independent experiments.

To determine if TMED9 is involved in selective autophagy, an mt-Keima assay was employed to determine mitophagy^57^. Upon carbonyl cyanide chlorophenylhydrazone (CCCP) treatment of HeLa cells expressing Parkin^58^, the portion of 561 nm excitable signal was increased in both flow cytometry and fluorescence imaging analyses, indicating the induction of mitophagy (Fig. 2G-J). The switch of mt-Keima excitation was partially blocked (∼40% decrease compared to CCCP treatment without TMED9 RNAi) in the absence of TMED9, suggesting the involvement of TMED9 in mitophagy (Fig. 2G-J). In addition, depletion of TMED9 also compromised the lysosome-dependent turnover of an aggregation-prone protein SOD1(G93A)^59^, suggesting that TMED9 may also regulate aggrephagy (Fig. 2K).

### TMED9 regulates membrane contact between ERGIC and ERES

Previously, we found that starvation induces enlargement of SEC12-positive ERES, which extends around the ERGIC^60^. To determine the effect of TMED9 depletion on ERES and ERGIC, we employed Stochastic Optical Reconstruction Microscopy (STORM) analysis (Fig. 3A, B). Consistent with our previous finding^60^, starvation induced formation of elongated ERES enwrapping the ERGIC (Fig. 3A). Notably, ERES and ERGIC were departed in the absence of TMED9 (Fig. 3A). Quantification shows that the association events between the ERGIC and ERES were reduced from ∼60 to ∼20 (∼3-fold decrease) in every 100 pairs of ERGIC-ERES structures analyzed (Fig. 3B). Using three-color STORM, we observed the distribution of TMED9 at the association sites of the ERGIC and ERES (Fig. 3C). Therefore, the data indicate that TMED9 regulates ERGIC-ERES association.

**Figure 3.**
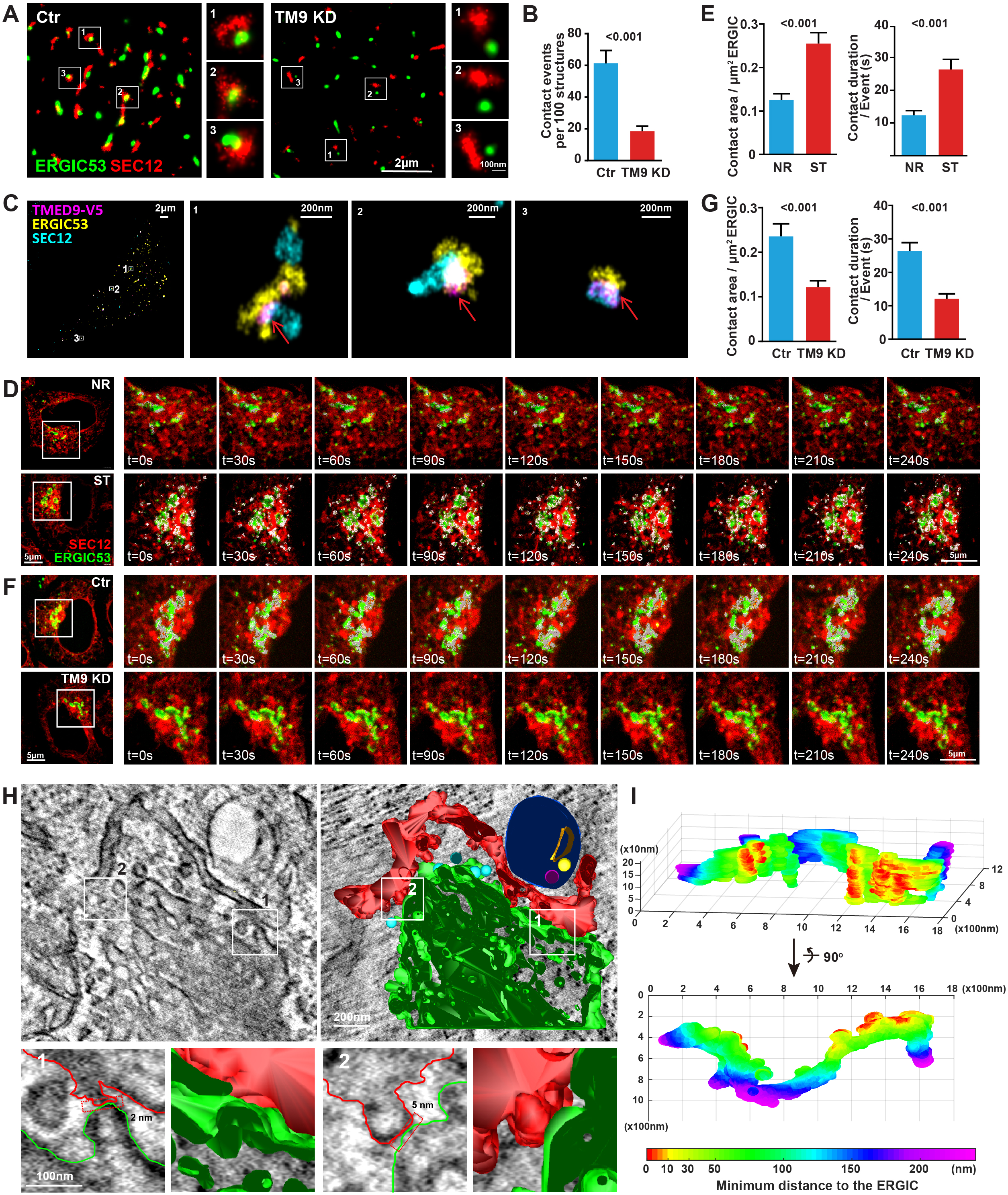
TMED9 modulates ERES-ERGIC contact. (A) 3D-STORM analysis of HeLa cells (control and TMED9 KD) labeled with anti-ERGIC-53 and SEC12 antibody. The cells were starved in EBSS for 1 h. (B) Quantification of contact events per 100 structures as shown in (A). Error bars represent standard deviations of > 20 cells from three independent experiments (> 6 cell per experiment). P-value was obtained from two-tailed t-test. (C) 3D-STORM analysis of HeLa cells expressing TMED9-V5 labeled with ERGIC-53 SEC12, and V5 antibodies. The cells were starved in EBSS for 1 h. Arrows point to the TMED9-enriched ERGIC-ERES associated region. (D, F) SD-SIM analysis of HeLa cells stably expressing EGFP-ERGIC-53 and mCherry-SEC12 incubated in nutrient-rich medium (NR) or starved (ST) in EBSS (D) or HeLa cells transfected with control or siRNAs against TMED9 starved in EBSS (F). Live imaging was performed starting from 30 min after the start of each treatment and pictures of each of the time points were shown. Contact analysis (white puncta) was performed using Imaris. (E, G) Quantification of contact area (μm^2^/100 μm^2^ ERGIC area) and contact duration in (D and F). Error bars represent standard deviations of > 30 cells from three independent experiments (> 10 cell per experiment). P-value was obtained from two-tailed t-test. (H) 3D-tomography of ERGIC-ERES in HeLa cells starved in EBSS for 1 h. The values shown in the insets represent two tight contact sites between the ERES and ERGIC. Scale bar sizes are indicated in the picture. Red: ERES; green: ERGIC; cyan: transport vesicles; blue: autophagosome; orange, yellow, and magenta: autophagic cargos. (I) Heatmap showing the distances between the ERES and ERGIC in (H). The results were calculated in and plotted using MATLAB. The X, Y, and Z axes show the dimensions of ERES (nm) in the tomogram. The scale bar represents the contact distance (nm).

We performed living-cell imaging using Spinning Disk Structured Illumination Microscopy (SD-SIM)^61^ to analyze the dynamics of ERGIC-ERES association (Fig. 3D-G, Video. S1-4). Under steady states, around 12% of the ERGIC surface (0.12 μm^2^ of contact per 1 μm^2^ ERGIC) associated with ERES labeled with SEC12 (Fig. 3D, E). The average life time of each point of association was around 12 seconds (Fig. 3E). Starvation increased the ERGIC-ERES association as measured by surface area and duration (∼2-fold increase in surface area, and ∼2.2-fold increase in time, Fig. 3D, E). TMED9 depletion reduced ERGIC-ERES association upon starvation, inducing a ∼2-fold decrease in surface area and ∼2.1-fold reduction in duration (Fig. 3F, G).

To determine if the ERGIC-ERES association is characterized by membrane contact, we performed 3D-tomography through EM. Indeed, ERGIC-ERES formed tight membrane contact sites along the associated portion of the two compartments (Fig. 3H, I). Within the contact sites, we observed tight apposition of the ERGIC-ERES membranes as narrow as 2-5 nm (Fig. 3H), and no obvious membrane fusion was observed at the contact sites. We quantified the minimal distance to the ERGIC from the ERES surface. In the heatmap based on the analyzed ERGIC-ERES contact, we observed contact regions (< 30 nm distant including both yellow and red) on the ERES, with a total surface area of ∼3.26 μm^2^, in which the area in close contact (< 10 nm, red) was ∼0.78 μm^2^ (Fig. 3I).

### TMED9 interacts with SEC12 via the C-terminal cytoplasmic tail

SEC12 was shown to concentrate in the ERES portion that associates with the ERGIC^60^. In co-immunoprecipitation analyses, SEC12 associated with TMED9, which was enhanced by starvation (Fig. 4A, B). Consistent with previous work^62^, TMED9 also formed a complex with TMED2 and TMED10 (Fig. 4A). Truncation analyses revealed that the C-terminal half of the SEC12 cytoplasmic domain associates with TMED9 (Fig. 4C). The C-terminal tail (CT) of TMED9 interacted with both the full length and the cytoplasmic domain of the SEC12 protein (Fig. 4D; Fig. S2A, B); Moreover, the CT of TMED9 is sufficient for SEC12 association (Fig. 4E). Furthermore, mutation of the FE residues to AA (m4) largely compromised the ability of TMED9-CT to associate with SEC12 (Fig. 4F). In an in vitro pull down experiment, TMED9-CT directly bound to the SEC12 cytoplasmic domain, which was abolished by the FE-AA mutation (Fig. 4G). Therefore, TMED9 directly interacts with the SEC12 cytoplasmic domain via the CT. The coil-coil (CC) domain of TMED9 is required for complex formation with full length SEC12, but not the SEC12 cytoplasmic domain, suggesting the involvement of TMED9 oligomerization in regulating TMED9-SEC12 association on the ERGIC-ERES (Fig. 4D; Fig. S2B).

**Figure 4.**
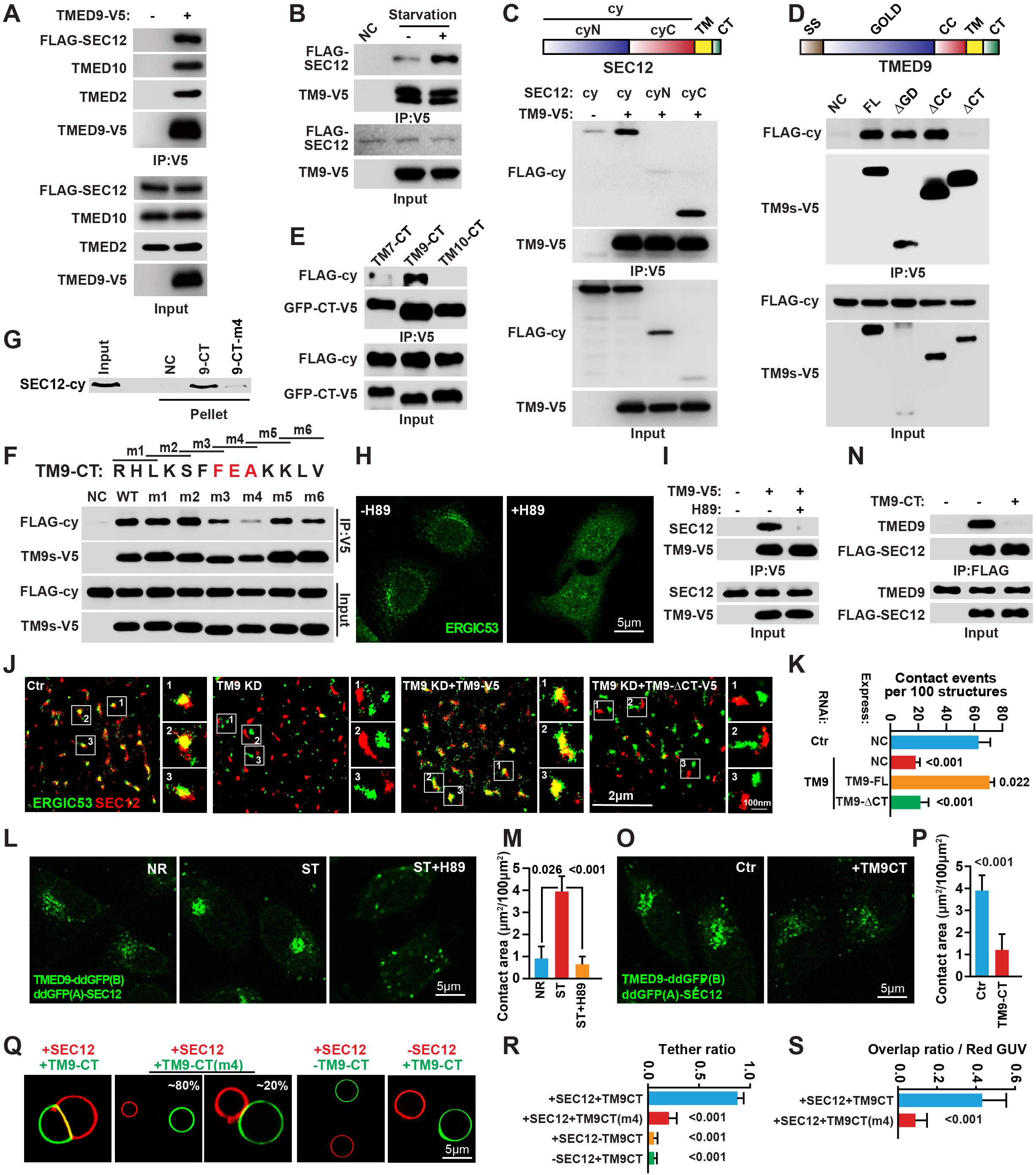
The interaction between TMED9 and SEC12 drives ERGIC-ERES contact formation. (A) Co-immunoprecipitation (CoIP) analysis of TMED9-V5 with FLAG-SEC12, TMED10, and TMED2 in HEK293T using anti-V5 agarose. (B) CoIP analysis of TMED9-V5 with FLAG-SEC12 in HEK293T incubated in nutrient-rich medium or starved in EBSS for 1 h using anti-V5 agarose. (C) CoIP analysis of FLAG-SEC12 variants (1-386 (cy); 1-239 (cyN) and 240-386 (cyC)) with TMED9-V5 in HEK293T starved in EBSS for 1 h using anti-V5 agarose. (D) CoIP analysis of TMED9-V5 variants (FL, full length TMED9; ΔGOLD, GOLD domain-deleted TMED9; ΔCC, CC domain-deleted TMED9 and ΔCT, C-terminal tail-deleted TMED9) with FLAG-SEC12 in HEK293T starved in EBSS for 1 h using anti-V5 agarose. (E) CoIP analysis of TMED7-CT, TMED9-CT, and TMED10-CT (triple repeats of CT with GFP in the N-terminus and V5 tag in the C-terminus) with FLAG-SEC12-cy in HEK293T starved in EBSS for 1 h using anti-V5 agarose. (F) CoIP analysis of TMED9-V5 variants (WT, full length TMED9; m1, R223A/H224A/L225A; m2, L225A/K226A/S227A; m3, S227A/F228A/F229A; m4, F229A/E230A; m5, K232A/K233A; m6, K233A/L234A/V235A) with FLAG-SEC12-cy in HEK293T starved in EBSS for 1 h using anti-V5 agarose. (G) Peptides (9-CT, CT of TMED9; 9-CT-m4, CT of TMED9-m4 in (F)) were immobilized on agarose beads followed by analysis of FLAG-SEC12 interaction in an in vitro pull-down assay. (H) Immunofluorescence of HeLa cells treated with 100 μM H89 for 20 min with ERGIC-53 antibody. (I) CoIP analysis of TMED9-V5 with FLAG-SEC12 in HEK293T treated with 100 μM H89 for 20 min using anti-V5 agarose. (J) Immunofluorescence and 3D-STORM of HeLa cells (control, TMED9 KD, TMED9 KD with TMED9-V5 or TMED9-ΔCT-V5 expression) with anti-ERGIC-53 and SEC12 antibody. The cells were starved in EBSS for 1 h. (K) Quantification of contact events per 100 structures as shown in (J). Error bars represent standard deviations of > 20 cells from three independent experiments (> 6 cells per experiment). P-value was obtained from two-tailed t-test. (L) Fluorescence images of HeLa cells stably expressing ddGFP(A)-SEC12 and TMED9-ddGFP(B) incubated in nutrient-rich medium or starved in EBSS for 1 h with and without 100 μM H89 treatment for 20 min. (M) Quantification of contact area (μm^2^/100 μm^2^ cell area) as shown in (L). Error bars represent standard deviations of > 150 cells from three independent experiments (> 50 cells per experiment). P-value was obtained from two-tailed t-test. (N) CoIP analysis of FLAG-SEC12 with TMED9 in HEK293T treated with Tat-TM9CT peptides (50 μM, 3 h) and starved in EBSS for 1 h using anti-FLAG agarose. (O) Fluorescence images of HeLa cells stably expressing ddGFP(A)-SEC12 and TMED9-ddGFP(B) treated with control or Tat-TM9CT peptides (50 μM, 3 h) and starved in EBSS for 1 h. (P) Quantification of contact area (μm^2^/100 μm^2^ cell area) as shown in (O). Error bars represent standard deviations of > 150 cells from three independent experiments (> 50 cells per experiment). P-value was obtained from two-tailed t-test. (Q) Fluorescence images of contact formation between “ERES-GUVs” and “ERGIC-GUVs” with indicated proteins or peptides. (R, S) Quantification of tether/contact ratio (ratio of ERES-GUV in contact with ERGIC-GUV (R)) and overlap ratio (overlap area to ERES-GUV area, (S)) as shown in (Q). Error bars represent standard deviations of > 150 GUVs from three independent experiments (> 50 GUVs per experiment). P-value was obtained from two-tailed t-test.

TMED9 was synthesized in the ER before reaching the ERGIC. Therefore, two modes of TMED9-SEC12 interaction may exist. One mode is intra-compartment interaction in which TMED9 associates with SEC12 in the ER. The other mode is inter-compartment interaction where ERGIC-TMED9 interacts with ERES-SEC12. The later is able to establish membrane contact between the ERGIC and ERES in theory. Interestingly, treatment of cells with H89, which diminished the ERGIC^27^ and led to the retention of TMED9 in the ER, abolished the TMED9-SEC12 interaction (Fig. 4H, I), suggesting that the localization of TMED9 to the ERGIC is required for TMED9-SEC12 interaction, which favors the possibility that TMED9-SEC12 associates in an inter-compartment manner and may regulate ERGIC-ERES contact.

### TMED9-SEC12 interaction drives ERGIC-ERES contact formation

To determine the involvement of TMED9-SEC12 interaction in ERGIC-ERES contact formation, TMED9 was depleted by RNAi followed by re-expression of full length or CT deleted TMED9 (ΔCT, SEC12 binding-deficient). In the STORM imaging analysis, the full length TMED9 rescued the ERGIC-ERES contact formation, rather than the ΔCT TMED9 (Fig. 4J, K). To facilitate ERGIC-ERES contact analysis, we employed a dimerization-dependent GFP (ddGFP) system^63^ by fusing each half of the GFP to the cytoplasmic sides of TMED9 and SEC12. The complemented GFP signal appeared in the perinuclear region characteristic of the ERGIC and was enhanced by starvation, recapitulating starvation-enhanced ERGIC-ERES contact formation (Fig. 4L, M). We employed the ddGFP assay to confirm the role of TMED9-SEC12 interaction in ERGIC-ERES contact. H89 treatment, which blocked TMED9-SEC12 interaction (Fig. 4I), abolished the GFP complementation signal (Fig. 4L, M). In addition, treatment of cells with a TMED9-CT peptide, which competed with the TMED9-SEC12 interaction (Fig. 4N), compromised the ERGIC-ERES contact by 4-fold in the ddGFP system (Fig. 4O, P). Therefore, TMED9-SEC12 interaction is required for ERGIC-ERES contact formation.

To test if the TMED9-SEC12 interaction is sufficient to drive membrane contact formation, we attached the cytoplasmic domain of SEC12 (“ERES-GUV”) and TMED9-CT (“ERGIC-GUV”) to giant unilamellar vesicles (GUV). Notably, GUVs with SEC12 and TMED9-CT, respectively, formed extensive contacts, in which ∼90% of GUVs with opposing proteins interacted and > 40% of the surface of the SEC12-positive red GUVs overlapped with the TMED9-CT-positive green GUVs (Fig. 4Q-S). The SEC12 binding-deficient TMED9-CT mutant (m4) decreased GUV tethering by ∼4-fold (measured by tether ratio, Fig. 4Q, R) and ∼8-fold (measured by overlap area, Fig. 4Q, S). Therefore, the data demonstrate that the specific interaction between the SEC12 cytoplasmic domain and TMED9-CT is sufficient to establish a membrane contact in vitro.

### TMED9-SEC12 interaction is required for autophagosome biogenesis

In an immunofluorescence assay, expression of wild type (WT) TMED9 rescued LC3 puncta formation in TMED9-deficient cells in response to starvation (Fig. 5A, B). The effect of restoration was reduced by ∼2-fold in TMED9-ΔCT or TMED9 m4 mutant indicating the requirement of TMED9-SEC12 interaction for regulating autophagosome biogenesis (Fig. 5A, B). Consistently, TMED9-ΔCT or m4 mutant compromised the restoration of LC3 lipidation mediated by WT TMED9 in TMED9-deficient cells (Fig. 5C). To further confirm the essential role of TMED9-SEC12 interaction in autophagosome biogenesis, overexpression of the TMED9-CT domain, which impairs TMED9-SEC12 interaction (Fig. 5D), was performed. Again, the TMED9-CT domain decreased autophagosome formation by ∼2-fold as measured by puncta formation of the endogenous LC3 (Fig. 5E, F), and largely abolished autophagosome and autolysosome formation in the tandem fluorescence LC3 assay (Fig, 5G-I; > 3-fold decrease). Therefore, ERES-ERGIC contact mediated by TMED9-SEC12 interaction is required for autophagosome biogenesis.

**Figure 5.**
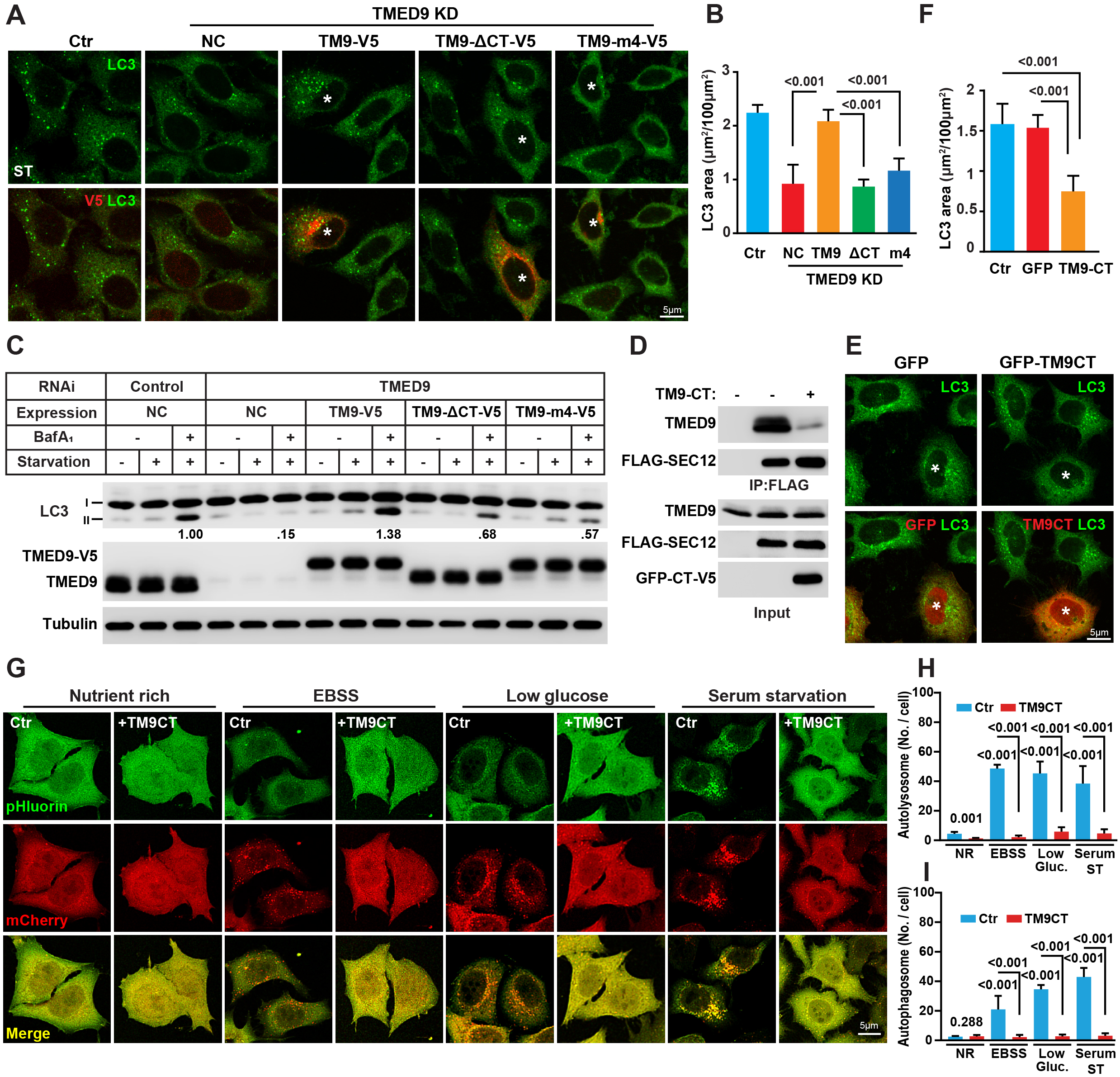
The interaction of TMED9 and SEC12 regulates autophagosome biogenesis. (A) Immunofluorescence of HeLa cells (control, TMED9 KD, TMED9 KD with TMED9-V5, TMED9-ΔCT-V5, or TMED9-m4-V5 expression) with V5 and LC3 antibodies. The cells were starved in EBSS for 1 h. Asterisks indicate cells with indicated TMED9 variant expression. (B) Quantification of the LC3 puncta area (μm^2^/100 μm^2^ cell area) analyzed in (A). Error bars represent standard deviations of > 150 cells from three independent experiments (> 50 cells per experiment). P-value obtained from two-tailed t-test. (C) LC3 lipidation in HeLa cells transfected with control or siRNAs against TMED9 with or without TMED9-V5, TMED9-ΔCT-V5, or TMED9-m4-V5 re-expression. The cells were incubated in nutrient-rich medium or starved in EBSS in the absence or presence of 500 nM bafilomycin A1 for 1 h. Immunoblots were performed to determine the levels of indicated proteins. Quantification was performed similarly to Fig. 1G. The blots are representative of at least three independent experiments. (D) CoIP analysis of FLAG-SEC12 with TMED9 in HEK293T transfected with or without GFP-tagged triple CT of TMED9 using anti-FLAG agarose. The cells were starved in EBSS for 1 h. (E) Immunofluorescence of HeLa cells transfected with GFP-tagged triple CT of TMED9 after EBSS starvation (1h) with anti-V5 and LC3 antibodies. Asterisks indicate cells with indicated protein expression. (F) Quantification of the LC3 puncta area (μm^2^/100 μm^2^ cell area) analyzed in (E). Error bars represent standard deviations of > 150 cells from three independent experiments (> 50 cells per experiment). P-value was obtained from two-tailed t-test. (G) HeLa cells stably expressing mCherry-pHluorin-LC3B were treated with control or Tat-TM9CT peptides (50 μM, 4 h) and incubated in nutrient-rich medium, EBSS, low glucose medium, or medium without serum. Confocal microscopy was performed. (H, I) Quantification of autolysosome (red, (H)) and autophagosome (yellow, (I)) numbers per cell as shown in (G). Error bars represent standard deviations of > 300 cells from three independent experiments (> 100 cells per experiment). P-value was obtained from two-tailed t-test.

### ERGIC-ERES contact facilitates SEC12 translocation to the ERGIC and COPII vesicle formation on the ERGIC

Previously, we detected an event of SEC12 translocation from ERES to ERGIC, which triggers the generation of a new type of COPII vesicle, termed the ERGIC-COPII vesicle^29,60^. Instead of the acting as a cargo carrier for ER-Golgi trafficking, the ERGIC-COPII vesicle fulfills the role of a lipidation precursor for autophagosome biogenesis^29^. To determine if ERGIC-ERES contact regulates COPII assembly on the ERGIC, the effect of TMED9 RNAi on the relocation of SEC12 to the ERGIC upon starvation was analyzed (Fig. 6A). Disrupting ERGIC-ERES contact by TMED9 RNAi abolished the relocation of SEC12 to the ERGIC (Fig. 6A, ∼5-fold decrease). Similarly, upon starvation, TMED9 depletion also decreased SEC31A (a COPII component) colocalization with the ERGIC by ∼3-fold (Fig. 6B, C). Therefore, ERGIC-ERES contact is required for SEC12 relocation to the ERGIC, which triggers assembly of COPII.

**Figure 6.**
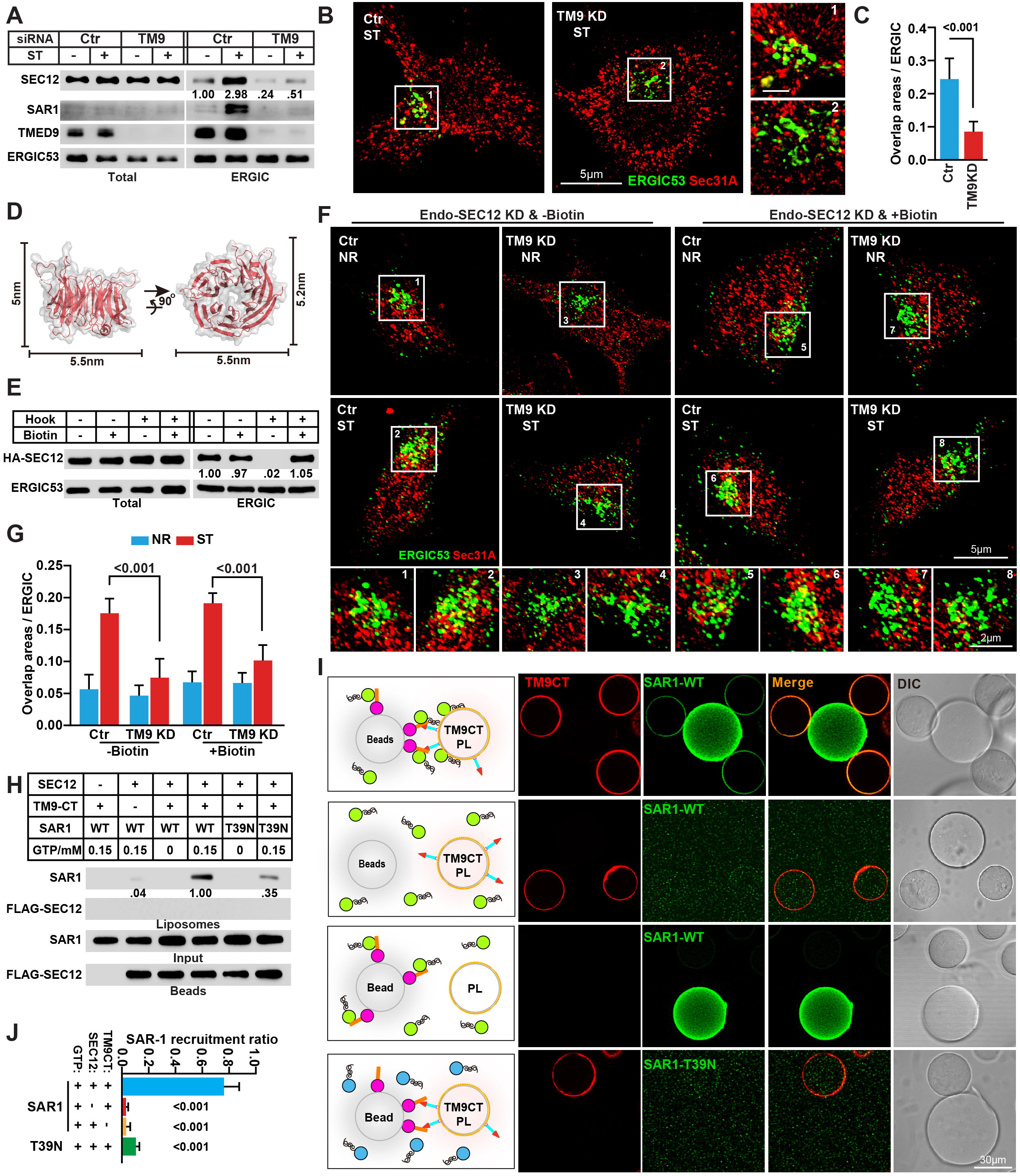
The function of ERGIC-ERES contact in ERGIC-COPII formation. (A) HEK293T cells were transfected with control or siRNAs against TMED9. After 72 h, the cells were incubated in nutrient-rich medium or starved in EBSS for 1 h. The cell lysates and the ERGIC membrane fractions were analyzed by immunoblot to determine the levels of indicated proteins. Quantification shows relative SEC12 relocation to the ERGIC under the indicated conditions. The control siRNA transfection group with nutrient-rich medium treatment was set as 1.00. The blots are representative of at least three independent experiments. (B) SIM analysis of ERGIC and COPII in HeLa control cells and TMED9 KD with ERGIC-53 and SEC31A antibodies. The cells were starved in EBSS for 1 h. (C) Quantification of COPII overlap area with the ERGIC (μm^2^/100 μm^2^ ERGIC area) as shown in (B). Error bars represent standard deviations of > 150 cells from three independent experiments (> 50 cell per experiment). P-value was obtained from two-tailed t-test. (D) The structure of the SEC12 cytoplasmic domain (PDB:5tf2). The values shown in the figure represent the dimension of SEC12. Structure model was created by PyMOL. (E) HEK293T cells were infected with RUSH-SEC12 lentivirus. The cells were treated with or without 40 μM biotin for 1 h. The cell lysates and the ERGIC membrane fractions were analyzed by immunoblot to determine the levels of indicated proteins. Quantification shows the percentage of SEC12 relocation to the ERGIC under the indicated conditions. (F) SIM analysis of ERGIC and COPII in HeLa cells stably expressing RUSH-SEC12. The cells were transfected with siRNAs against SEC12 and TMED9, incubated in nutrient-rich medium or EBSS for 1 h with or without 40 μM biotin for 1 h, and labeled with ERGIC-53 and SEC31A antibodies. (G) Quantification of COPII overlap area with the ERGIC (μm^2^/100 μm^2^ ERGIC area) as shown in (F). Error bars represent standard deviations of > 150 cells from three independent experiments (> 50 cells per experiment). P-value was obtained from two-tailed t-test. (H) Immunoblot showing the transactivation of SAR1 on the liposome via SEC12-bound beads. Beads with or without SEC12 were incubated with liposomes with or without TMED9-CT, together with indicated variants of SAR1 in the presence or absence of GTP. After reaction, the liposomes were isolated followed by immunoblot. Quantification was based on the ratio of SAR1 with the group of highest transactivation (with SEC12, TM9-CT, SAR1, and 0.15 nM GTP set as 1.00). The blots are representative of at least three independent experiments. (I) Fluorescence imaging showing the recruitment of SAR1-BFP to the TMED9-CT-labeled liposomes attached to streptavidin agarose beads. Beads with or without SEC12 were incubated with beads coated with control liposomes or TMED9-CT. Indicated SAR1-BFP variants with GTP were incubated with the indicated combination of beads. Confocal imaging was performed to analyze the recruitment of SAR1-BFP to the liposomes on the beads in contact with beads with SEC12. (J) Quantification of SAR1 recruitment ratio as shown in (I). Error bars represent standard deviations of > 150 beads with liposomes from three independent experiments (> 50 beads with liposomes per experiment). P-value obtained from two-tailed t-test.

### ERGIC-ERES contact allows transactivation of ERGIC-COPII vesicle formation by ERES-localized SEC12

As described above, the ERGIC-ERES forms tight contact with distances as short as 2-5 nm (Fig. 3H, I). The structure of the SEC12 cytoplasmic domain (PDB:5tf2) was revealed to have dimensions of 5.5 × 5 × 5.2 nm (Fig. 6D). Therefore it is possible that the ERES-localized SEC12 reaches the ERGIC membrane in the contact region. Considering the ERES harbors the majority of SEC12, we questioned if ERES-localized SEC12 is able to transactivate COPII vesicle formation in the ERGIC. To test this possibility, we employed a Retention Using Selective Hooks (RUSH) system to retain SEC12 on the ER^64^. Indeed, The Streptavidin-Binding Peptide (SBP)-tagged SEC12 failed to translocate to the ERGIC in the presence of the hook protein (major histocompatibility complex (MHC) class II-associated invariant chain (Ii) fused with streptavidin) (Fig. 6E). The addition of biotin, which released SBP-SEC12 from the hook, allowed the translocation of SEC12 to the ERGIC upon starvation (Fig. 6E).

To determine the specific effect of SBP-SEC12 retention in the ERES, we depleted endogenous SEC12 by RNAi and expressed SBP-SEC12 with the hook. Interestingly, prevention of SBP-SEC12 translocation to the ERGIC only partially reduced autophagosome biogenesis (∼30% and ∼25% reduction in autophagosome and autolysosome production relative to those with biotin treatment, Fig. S3A, B) compared to TMED9 RNAi (∼80% decrease in both autophagosome and autolysosome production, Fig. S3A, B). The partial reduction of autophagosome biogenesis was also confirmed by analyzing LC3 lipidation, in which we observed a 23% decrease of LC3 lipidation (with starvation and bafilomycin A1) by retaining SEC12 on the ERES, compared to a ∼70% decrease by means of TMED9 RNAi (Fig. S3C). Accordantly, retaining SEC12 on the ERES only partially reduced COPII assembly on the ERGIC (Fig. 6F, G; ∼11% decrease in SEC31A colocalization with the ERGIC) compared to TMED9 RNAi (∼50% decrease in SEC31A colocalization with the ERGIC). The data support the notion that SEC12 translocation to the ERGIC is not the sole mechanism for TMED9-regulated ERGIC-COPII assembly and autophagosome biogenesis. Therefore, it is likely that the concentrated SEC12 on the ERES may transactivate COPII assembly on the ERGIC via the ERGIC-ERES contact.

To test the possibility of SEC12 transactivation of COPII assembly on the apposed membrane, we employed two in vitro approaches based on SAR1 GTP loading which is promoted by SEC12 GTP exchange factor activity and leads to the association of SAR1 with the membrane^65^. In one experiment, we attached TMED9-CT to liposomes, linked SEC12 cytoplasmic domain to agarose beads, and incubated them with SAR1 and GTP. After incubation, we analyzed the liposome-associated SAR1 as a readout of SAR1 transactivation by the SEC12 on the beads. As shown in Figure 6H, the presence of TMED9-CT in the liposome together with SEC12 on the beads, led to the presence of SAR1 in the liposome fraction, which was not observed without TMED9-CT, SEC12, or GTP, indicating the transactivation of SAR1 in the liposome by the SEC12 on the beads (Fig. 6H). The GTP binding-deficient SAR1 mutant (T39N) showed compromised localization (∼3-fold decrease) to the liposome, further confirming that the GTP binding catalyzed by SEC12 accounts for the SAR1 membrane association (Fig. 6H). In all tested groups, SEC12 was detected on the beads but not in the liposome fraction, confirming the effect of transactivation (Fig. 6H). In the other experiment, we coated streptavidin agarose beads with TMED9-CT labeled liposomes based on a previous study^66^, crosslinked SEC12 to agarose beads and incubated the beads with SAR1-BFP and GTP. In the fluorescence imaging assay, the presence of both TMED9-CT-liposomes and SEC12 on opposing beads promoted the recruitment of SAR1-BFP to the TMED9-CT-localized liposomes on the beads when they came into contact (Fig. 6I, J). Again the SAR1 mutant (T39N) was deficient when recruiting to TMED9-CT-localized beads (Fig. 6I, J; ∼80% decrease compared to WT SAR1). Together, the data indicate that SEC12 is able to transactivate SAR1 on an apposed membrane when contact mediated by TMED9-CT-SEC12 interaction is formed.

### Starvation-induced ERES enlargement is required for ERGIC-ERES contact formation

Previously, we found the enlargement of ERES upon starvation, which was mediated by SEC12-FIP200 association and maintained by CTAGE5^60^. We also determined the relationship between ERES remodeling and ERGIC-ERES contact formation. Consistent with that previous work, knockdown of CTAGE5 completely dispersed the ERES, whereas FIP200 depletion did not affect steady state ERES morphology but prevented starvation-induced ERES enlargement (Fig. 7A, B). Disruption of ERES enlargement by CTAGE5 or FIP200 RNAi decreased the TMED9-SEC12 interaction, as revealed by co-immunoprecipitation (Fig. 7C, D). FIP200 or CTAGE5 RNAi abolished ERGIC-ERES contact as exhibited by both STORM and ddGFP assays (Fig. 7E-H). The data indicate that enlargement of the ERES is required for TMED9-SEC12 interaction and ERGIC-ERES contact formation. On the contrary, knockdown of TMED9 moderately increased SEC12-FIP200 association (Fig. 7I, ∼1.2-fold increase) and consistently increased steady state ERES size (∼1.5-fold increase), which was not further increased upon starvation (Fig. 7A, B). The enlarged ERES at steady state caused by TMED9 RNAi was reversed by co-knockdown of CTAGE5 or FIP200 (Fig. 7A, B). The data suggest that ERES-remodeling occurs upstream of ERGIC-ERES contact formation under starvation conditions and, on the other hand, contact formation may also control steady state ERES size regulation by FIP200 as a feedback.

**Figure 7.**
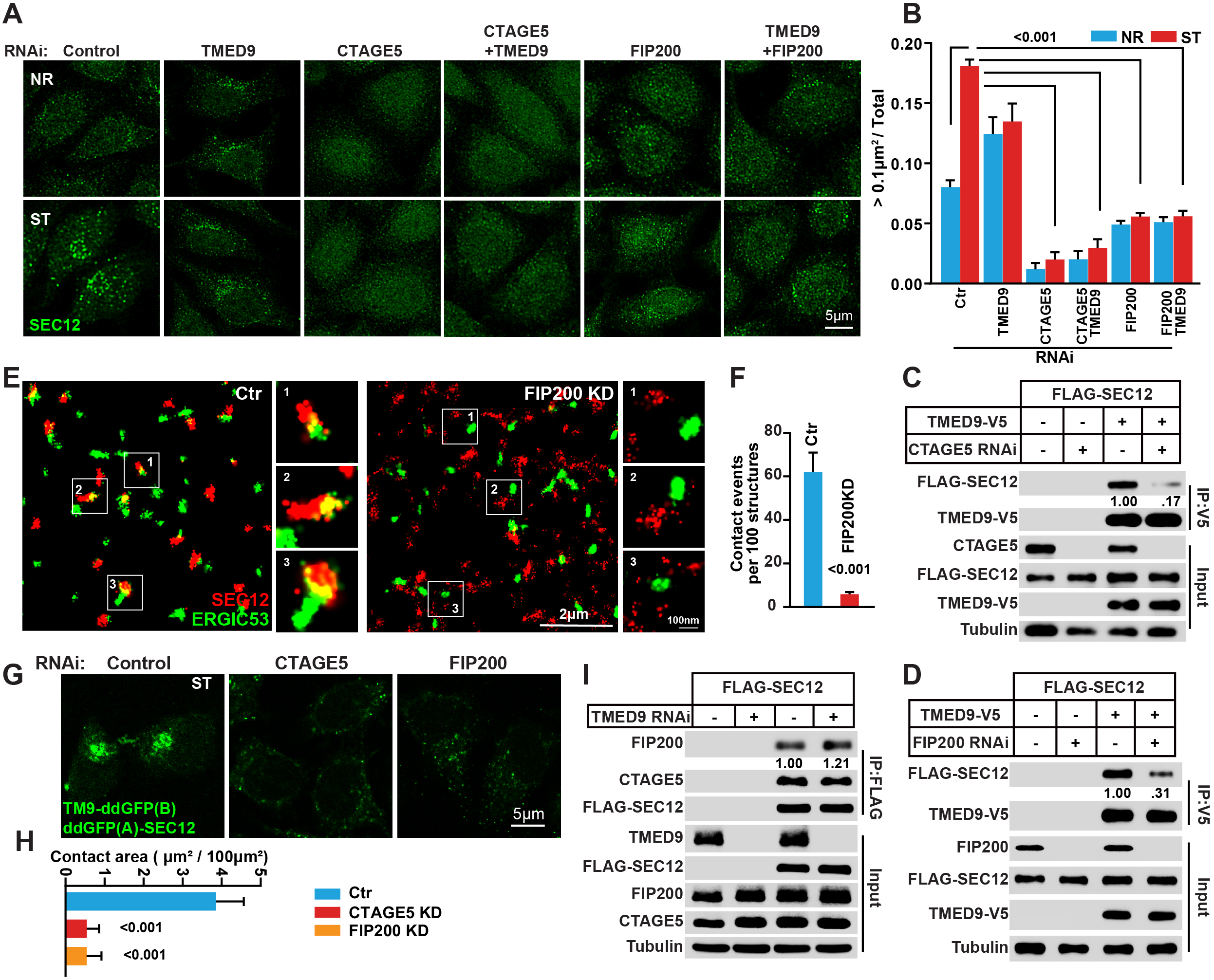
The requirement of ERES enlargement in ERGIC-ERES contact formation. (A) Immunofluorescence of HeLa cells transfected with control and siRNAs against TMED9, CTAGE5, or FIP200 with the SEC12 antibody. The cells were incubated in nutrient-rich medium or starved in EBSS for 1 h. (B) Quantification of the SEC12 puncta area (ratio of puncta > 0.1μm^2^) analyzed in (A). Error bars represent standard deviations of > 150 cells from three independent experiments (> 50 cells per experiment). P-value was obtained from two-tailed t-test. (C, D) CoIP analysis of TMED9-V5 with FLAG-SEC12 in HEK293T transfected with control or siRNAs against CTAGE5 (C) or FIP200 (D) using anti-V5 agarose. The cells were starved in EBSS for 1 h. Quantification was based on the ratio of FLAG-SEC12 to TMED9-V5 of IP fraction with the control setting as 1.00 (control siRNA). The blots are representative of at least three independent experiments. (E) 3D-STORM analysis of HeLa cells transfected with control and siRNAs against FIP200 with anti-ERGIC-53 and anti-SEC12 antibodies. The cells were starved in EBSS for 1 h. (F) Quantification of contact events per 100 structures as shown in (E). Error bars represent standard deviations of > 20 cells from three independent experiments (> 6 cells per experiment). P-value obtained from two-tailed t-test. (G) Fluorescence images of HeLa cells stably expressing ddGFP(A)-SEC12 and TMED9-ddGFP(B) transfected with control and siRNAs against CTAGE5 or FIP200. The cells were starved in EBSS for 1 h. (H) Quantification of contact area (μm^2^/100 μm^2^ cell area) as shown in (G). Error bars represent standard deviations of > 150 cells from three independent experiments (> 50 cells per experiment). P-value obtained from two-tailed t-test. (I) CoIP analysis of FLAG-SEC12 with FIP200 and CTAGE5 in HEK293T transfected with control and siRNAs against TMED9 using anti-FLAG agarose. The cells were starved in EBSS for 1 h. Quantification was based on the ratio of FIP200 to FLAG-SEC12 of IP fraction with a control setting of 1.00 (control siRNA). The blots are representative of at least three independent experiments.

### The ERGIC-ERES contact is distinct from TFG-mediated ERES-ERGIC tethering

Trk-fused gene (TFG) was recently shown to oligomerize and maintain the close position of ERES and ERGIC as well as to control the direction of COPII vesicle targeting^41–43^. To determine the relationship between TFG-regulated ERES-ERGIC clustering and ERES-ERGIC contact, a comparison of TFG knockdown with TMED9 knockdown was performed. As shown in Figure 8A, knockdown of TFG decreased the secretion of ssGFP, which echoes the effect of TFG on ER-derived COPII vesicle transport, whereas TMED9 RNAi did not affect ssGFP secretion (Fig. 8A). TMED9 depletion abolished autophagosome biogenesis as revealed by LC3 lipidation and the tandem-fluorescence LC3 assay (Fig. 8B & Fig. 2D-F). TFG knockdown moderately inhibited lipidated LC3 accumulation in the presence of bafilomycin A1 (∼14% decrease, Fig. 8B), which is consistent with a recent study showing the involvement of a later step of autophagy^67^. However, TFG knockdown did not affect autophagosome biogenesis or maturation, as revealed by the tandem-fluorescence LC3 assay (Fig. 8C-E). In the same line, TFG RNAi did not affect starvation-induced ERES enlargement, TMED9-SEC12 interaction, or ERGIC-ERES contact (Fig. 8F-L). Therefore, TFG and TMED9 function respectively in ER-Golgi trafficking and autophagosome biogenesis. Super-resolution imaging STORM demonstrated that TFG and TMED9 localized in distinct regions in the cell with few overlaps (Fig. 8M). In addition, the ddGFP signal generated by the ERGIC-ERES contact localized distinctly from TFG (Fig. 8N, O). Together the data indicate the TFG-mediated ERES-ERGIC cluster and TMED9-SEC12-established ERGIC-ERES contact are physically and functionally distinct in the ER-Golgi trafficking system.

**Figure 8.**
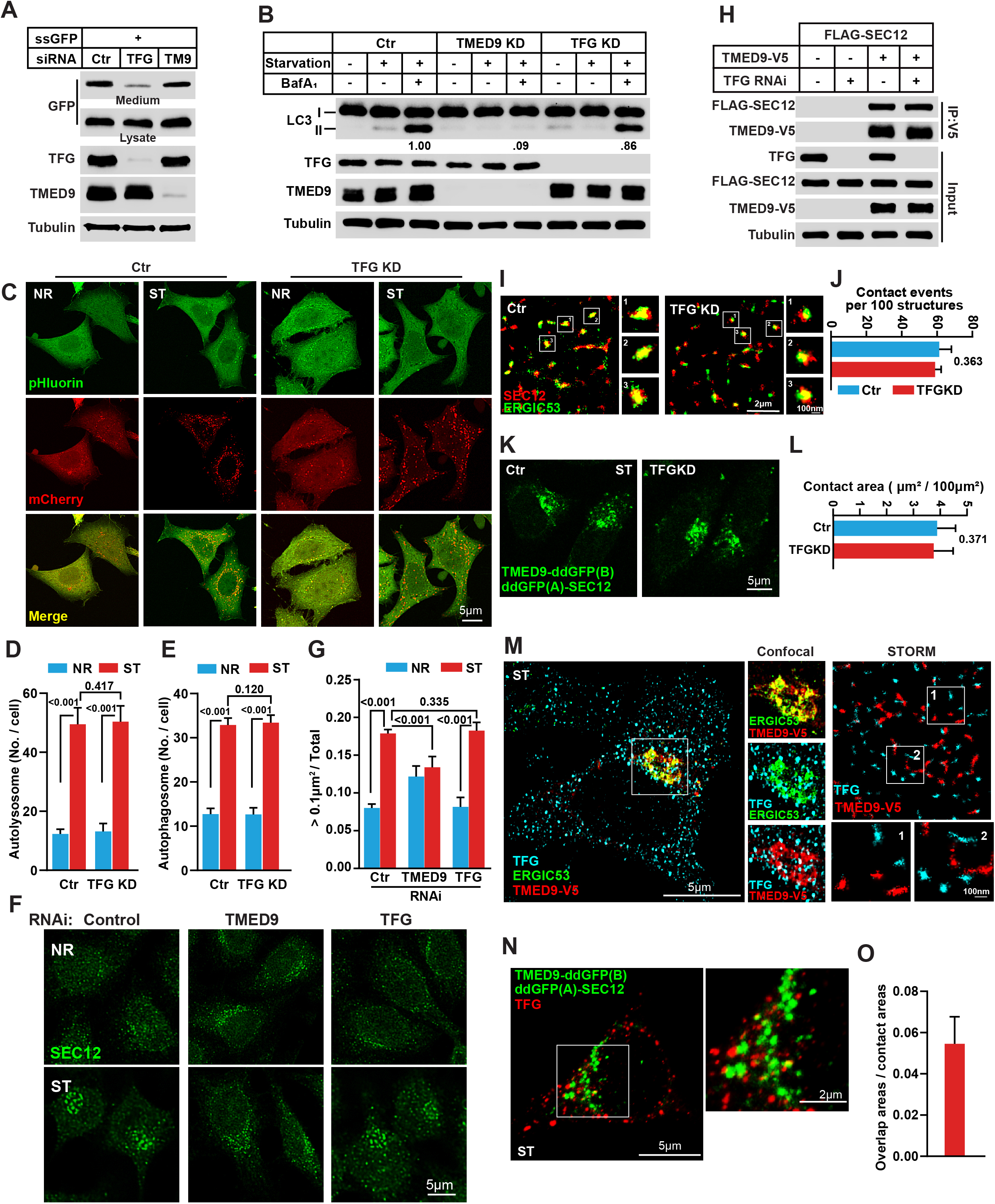
ERGIC-ERES contact is distinct from TFG-mediated ERES-ERGIC tethering. (A) Secretion of ssGFP in HEK293T transfected with control and siRNAs against TFG or TMED9. The blots are representative of at least three independent experiments. (B) LC3 lipidation in HeLa cells transfected with control or siRNAs against TMED9 or TFG. The cells were incubated in nutrient-rich medium or EBSS for 1 h in the absence or presence of 500 nM bafilomycin A1 for 1 h. Immunoblots were performed to determine the levels of indicated proteins. Quantification was performed similarly to Fig. 1G. The blots are representative of at least three independent experiments. (C) HeLa cells stably expressing mCherry-pHluorin-LC3B were transfected with control or siRNAs against TFG and incubated in nutrient-rich medium or starved in EBSS for 1 h. Confocal microscopy was performed. (D, E) Quantification of autolysosome (D) (red, from C) and autophagosome (E) (yellow, from E) numbers per cell as shown in (C). Error bars represent standard deviations of > 300 cells from three independent experiments (> 100 cell per experiment). P-values were obtained from two-tailed t-test. (F) Immunofluorescence of HeLa cells transfected with control or siRNAs against TMED9 or TFG with anti-SEC12 antibody. The cells were incubated in nutrient-rich medium or starved in EBSS for 1 h. (G) Quantification of the SEC12 puncta area (ratio of puncta >0.1 μm^2^) analyzed in (F). Error bars represent standard deviations of > 150 cells from three independent experiments (> 50 cells per experiment). P-value was obtained from two-tailed t-test. (H) CoIP analysis of TMED9-V5 with FLAG-SEC12 in HEK293T transfected with control and siRNA against TFG using anti-V5 agarose. The cells were starved in EBSS for 1 h. The blots are representative of at least three independent experiments (I) 3D-STORM analysis of HeLa cells transfected with control or siRNAs against TFG with anti-ERGIC53 and SEC12 antibodies. The cells were starved in EBSS for 1 h. (J) Quantification of contact events per 100 structures as shown in (I). Error bars represent standard deviations of > 20 cells from three independent experiments (> 6 cells per experiment). P-value was obtained from two-tailed t-test. (K) Fluorescence images of HeLa cells stably expressing ddGFP(A)-SEC12 and TMED9-ddGFP(B) transfected with control and siRNAs against TFG. The cells were starved in EBSS for 1 h. (L) Quantification of contact area (μm^2^/100 μm^2^ cell area) as shown in (K). Error bars represent standard deviations of > 150 cells from three independent experiments (> 50 cells per experiment). P-value was obtained from two-tailed t-test. (M) Immunofluorescence and 3D-STORM of HeLa cells transfected with TMED9-V5 with V5, ERGIC-53, and TFG antibodies. The cells were starved in EBSS for 1 h. (N) SIM analysis of HeLa cells stably expressing ddGFP(A)-SEC12 and TMED9-ddGFP(B) with anti-TFG antibody. The cells were starved in EBSS for 1 h. (O) Quantification of overlap area ratio (relative to ddGFP signal area) as shown in (N). Error bars represent standard deviations of > 150 cells from three independent experiments (> 50 cells per experiment).

## DISSCUSSION

Formation of the unique double-membrane autophagosome is a central step of autophagy and requires endomembrane reorganization to generate autophagosome precursors. The process of endomembrane remodeling has been a longstanding question awaiting a clear answer^4^. Here we identify a new membrane contact formed between the ERGIC and ERES, regulating the biogenesis of autophagosomal membrane precursors for LC3 lipidation. The interaction between TMED9 on the ERGIC and SEC12 on the ERES establishes ERGIC-ERES contact. Starvation triggers ERES-enlargement regulated by FIP200 and CTAGE5, as well as a TMED9-SEC12 interaction, which facilitates the formation of the ERGIC-ERES contact. The close contact (2-5 nm) between the ERGIC and ERES allows for translocation of SEC12 to the ERGIC to trigger ERGIC-COPII vesicle formation. Meanwhile the ERES-localized SEC12 transactivates COPII assembly on the ERGIC. Through the two mechanisms, the ERGIC-COPII vesicles for autophagosome biogenesis are generated during stress conditions (Fig. 9).

**Figure 9.**
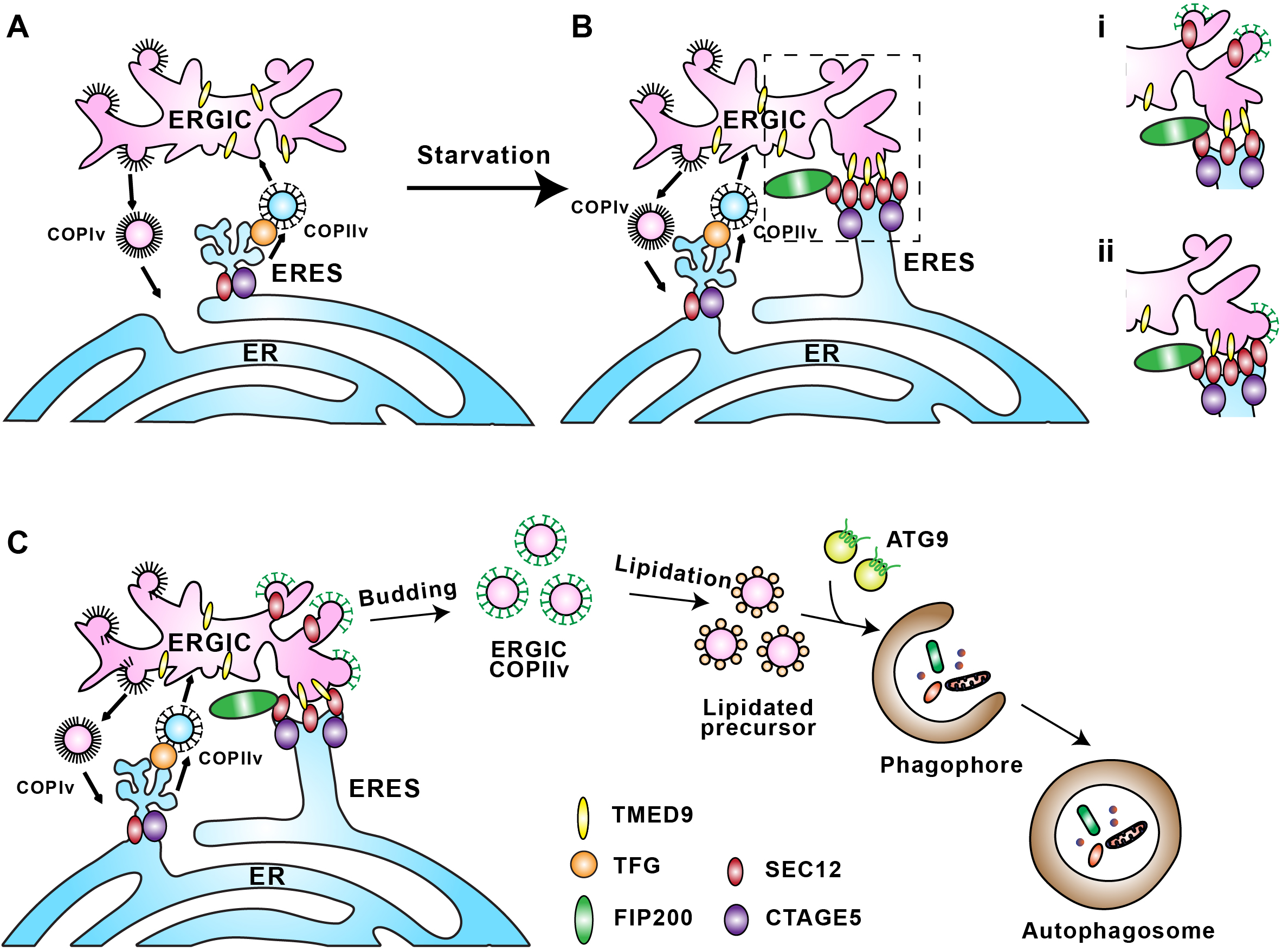
A model for ERES-ERGIC contact formation and autophagosome biogenesis. (A) In steady-state conditions, SEC12 (dark red oval) associates with CTAGE5 (purple oval) and localizes to the ERES. TFG (orange oval) tethers ERES and COPII vesicles for protein cargo transport to the ERGIC and Golgi. (B) Upon starvation, SEC12-ERES is enlarged and surrounds the ERGIC, the process of which is dependent on FIP200 (green oval) and CTAGE5. The remodeled ERES forms a contact with the ERGIC independent of TFG, which leads to the relocation of SEC12 to the ERGIC (i) and/or transactivation of COPII vesicle formation on the ERGIC by ERES-localized SEC12 (ii). (C) The ERES-ERGIC contact triggers the assembly of ERGIC-COPII vesicles as a membrane template for LC3 lipidation, a potential vesicular pool for the assembly of the pre-autophagosomal membrane.

Interestingly, depletion of TMED9 or CTAGE5 (in our previous work) preferentially abolished LC3 puncta formation instead of PAS assembly as revealed by FIP200 puncta (Fig. S1H, I)^60^. The data suggest that generation of autophagic membrane precursors for LC3 lipidation and the initiation of PAS assembly may be independently regulated. Nonetheless, autophagic factors, e.g. FIP200 and the class III PI3K, may regulate both processes, as depletion of these factors affects PAS formation and simultaneously inhibits multiple steps in ERGIC-ERES membrane remodeling upon starvation ^4,27,60^. The independent regulation of the two processes by the ATG proteins may be coordinated by differences in complex formation; e.g. FIP200 forms a complex with ULK1/ATG13/ATG101 at the PAS^4^, whereas it forms a complex with SEC12 in starvation-induced ERES remodeling^60^. It has been shown that the ERGIC-ERES system localizes adjacent to the PAS in both mammalian and yeast cells^15,68,69^. The proximity of the two sites may allow for efficient transfer and targeting of autophagic COPII vesicles to the PAS. In yeast, Ypt1 (the mammalian homologue of RAB1) and TRAPPIII were shown to regulate the delivery of COPII vesicles to the PAS^70–73^. The involvement of the ERGIC-localized RABs, RAB1 and RAB2, and TRAPPIII in autophagy has also been reported in mammalian cells^74–76^. However, clarification of whether the RABs, TRAPPIII, or other protein factors are involved in targeting the ERGIC-COPII vesicles to the PAS is unresolved.

Our data indicate that the ERGIC-ERES contact is a new type of compartment coordination with distinct structure and function in the ER-Golgi system. First, it is physically and functionally different from the previously known type of ERGIC-ERES association mediated by the oligomerized TFG. It has been shown that in the ERES, TFG associates with the ERES organizer SEC16, and the COPII inner coat component SEC23, to maintain the juxtaposed position of ERGIC-ERES, which is important for controlling the direction of COPII vesicle targeting and the efficiency of the transport^41–43^. In the case of ERGIC-ERES contact, the maintenance requires TMED9-SEC12 association but not TFG (Figs. 4, 8). Additionally, the formation of ERGIC-ERES contact is induced by starvation, and its major function is to regulate autophagosome biogenesis instead of protein transport in the secretory pathway (Figs. 3, 5, 8). Similarly, the time duration of ERES-cis-Golgi association was also increased under starvation conditions in a previous work using the yeast *Sacchoromyces cerevisiae*^77^. Considering the ERES has been shown to associate with the site of autophagosome biogenesis^68,69^, it is enticing to propose that formation of the ERGIC/Cis-Golgi-ERES contact for autophagosome biogenesis may be a conserved mechanism from yeast to mammals. Second, the ERGIC-ERES structure and COPII component distribution is distinct from those for the secretory pathway. Several works using different electron microscopical approaches dissected the structure of the ERES for ER-Golgi trafficking, and indicated a tubulovesicular clustered shape for the ERES^78–80^. Furthermore, two new studies implicated that several COPII components reside at the root of the ERES where the ER is connected^78,79^. Interestingly, the ERES involved in ERGIC-ERES contact is flatter (compared to the reported tubulovesicular structure of ERES), most likely to accommodate for the contact formation (Fig. 3H, I). In addition, ERGIC-ERES contact is facilitated by the TMED9-SEC12 association, implying the location of SEC12 on the distal side of the ERES in connection with the ERGIC.

One distinguishing feature of the ERGIC-ERES contact is its short distance (as close as 2-5 nm) compared to the conventional membrane contact of 10-30 nm between apposed organelles, e.g. the ER-mitochondria contact. This allows the SEC12 cytoplasmic domain (5.5×5×5.2 nm) to physically contact the ERGIC membrane triggering COPII formation on the ERGIC through transactivation. A similar example of transactivation was also reported in another type of membrane contact, the ER-trans-Golgi contact, in the ER-Golgi system ^81^. In that membrane contact, the distance is as close as 5 nm, and the ER-localized SAC1 was proposed to hydrolyze the PI4P on the Golgi in trans^82,83^. In addition to trans-activation, SEC12 also relocates from the ERES to the ERGIC as another mechanism for generating ERGIC-COPII vesicles. The contact synchronizes the action of the ERGIC and ERES through both mechanisms, which allows them to cooperate to generate the autophagosome precursors. How SEC12 is translocated to the ERGIC is pending. The process is unlikely to be mediated by membrane fusion because we did not observe any fusion events between the ERGIC-ERES contact surface.

The ERGIC was initially identified as a sorting station for membrane trafficking between the ER and the Golgi apparatus^84^. Notably, the current work, together with our previous studies, indicates that the ERGIC is a multi-functional compartment. In response to stress, the ERGIC acts as a membrane platform that orchestrates the generation of autophagosomes through contact formation with the ERES, and subsequently generates ERGIC-COPII vesicles. In addition, our recent work identified the ERGIC as a vesicle carrier for the translocation of secretory cargoes without a signal peptide for unconventional protein secretion^85^. Numerous studies have also found that the ERGIC is a membrane station supporting coronavirus assembly^86^. Even with respect to ER-Golgi trafficking, the ERGIC was shown to be sub-classified. A recent work found a tubular form of the ERGIC that allows for expedited travelling of SURF4 cargoes, which is distinct from canonical tubulovesicular ERGIC positive for ERGIC-53^87^. Therefore, the ERGIC could be classified as a dynamic membrane, compartment accommodating long distance travel between the ER and Golgi apparatus in mammalian cells, as well as coping with stress-related endomembrane remodeling events, which could be targeted by pathogens, including coronaviruses. It is likely that the ERGIC sub-compartmentalizes to meet multifunctional needs. Future work is necessary to understand how the ERGIC is subdivided, and what controls its subdivision.

## Acknowledgments

Dr. Ge is deeply grateful to Dr. Randy Schekman for the post-doctoral training at UC Berkeley. We thank Dr. Hongguang Xia (Zhejiang University, China), Dr. Houchaima Ben-Tekaya (University of Basel, Switzerland), Dr. Yueguang Rong (Huazhong University of Science and Technology, China) for reagents, Peng Zhao and Huijiao Bai at Tsinghua University for technical assistance. We thank Dr. Li Yu, Dr. Ye-Guang Chen (Tsinghua University, China), Dr. Wei Liu (Zhejiang University, China), Dr. Quan Chen (Nankai University, China), Dr. Qing Zhong (Shanghai Jiao Tong University, China) for helpful suggestions on the study. The authors would like to acknowledge the assistance of Imaging Core Facility, Technology Center for Protein Sciences, Tsinghua University.

## Author contributions

S. L. and G. L. performed experiments, analyzed data, and wrote the manuscript. R. Y., J. X., S. Z., X. M., Y. L., and J. G. L. performed experiments. M. Z., Q. S., L. C., S. L. and K. X. analyzed data.

## MATERIALS AND METHODS

### Plasmids and siRNAs

The TMEDs family members (TMED7, TMED9 and TMED10 were purchased from DNASU, and the others were amplified from HEK293T cDNA) were PCR amplified and inserted into the FUGW vector with a V5 tag at the C-terminus. The TMED9 truncations and mutations were generated by mutagenesis PCR. N-terminal EGFP-tagged ERGIC53 (from Dr. Houchaima Ben-Tekaya) was inserted into the FUGW vector. Human SEC12 was PCR amplified from Flag-SEC12 ^60^ and inserted into the FUGW vector with a mCherry tag at the N-terminus. N-terminal HA-tagged SEC12 domains (1-386, 1-239, and 240-386 aa) were described previously^60^. The MitoKeima plasmid was a gift from Dr. Hongguang Xia (Zhejiang University). The C-terminal BFP-tagged hSAR1a-WT and hSAR1a-T39N were inserted into PGEX4T1 vector for protein purification. The tandem fluorescence (mCherry-pHluorin) LC3 was a gift from Dr. Yueguang Rong (Huazhong University of Science and technology).

The siRNAs that were used in this work are listed in Table S2. An equimolar mixture of different siRNAs for a specific gene was used to induce gene silencing. AllStars negative siRNA (GenePharma) was used as a control. The siRNA transfection was performed with Lipofectamine RNAiMAX (Invitrogen, 13778150) according to the manufacture’s protocols.

### Antibodies, reagents and peptides

The antibodies that used in this work are listed in Table S2. The following reagents were purchased from the indicated sources: CCCP (Selleck, S6494), Cycloheximide (CST, 2112S), Bafilomycin A1 (Selleck, S1413), Rapamycin (Sigma, R8781), H89 (CST, 9844). Other reagents were described previously^60,85^.

L-amino acid peptides were synthesized by Scilight Biotechnology LLC. The Tat-TM9CT peptide sequence, YGRKKRRQRRRGGRHLKSFFEAKKLV, consisted of 11 amino acids from the Tat PTD at the N terminus, a GG linker to increase flexibility, and at the C terminus, 13 amino acids derived from TMED9 (223-235). The control peptide (Tat-scrambled) consisted of the Tat protein transduction domain, a GG linker, and a scramble sequence (YGRKKRRQRRRGGVGNDFFINHETTGFATEW)^88^. The FITC tagged TM9CT (FITC-TM9CT) peptide consisted of the FITC, an aminohexanoic acid, a Cysteine (for the binding with PE-MPB) and 15 amino acids derived from TMED9 (221-235) (QMRHLKSFFEAKKLV). The FITC-tagged TM9CT (m4) (FITC-TM9CT(m4)) peptide consisted of the FITC, an aminohexanoic acid, a Cysteine and 15 amino acids derived from TMED9 (221-235) containing two substitutions, including F229A and E230A (QMRHLKSFAAAKKLV). For peptide treatment, cells were washed with PBS and treated with peptides (40 μM, 1–4 h) dissolved in OPTI-MEM (Gibco) acidified with 0.15% (v/v) 6N HCl.

### Cell culture and transfection

Maintenance of cell lines and transfection were described previously^60,85^. Cells (HEK293T, U2OS, Hela) were maintained in DMEM supplemented with 10% FBS at 37°C in 5% CO_2_. Transfection was performed using PEI (Polysciences, Inc.) for HEK293T, PolyJet (Signagen, SL100688) for Hela and X-tremeGENE HP (Roche, 28088300) for U2OS according to the manufacture’s protocols.

### Lentiviral transduction and generation of CRISPR KO cells

For lentiviral transduction, HEK293T cells were transfected with pLX304 plasmids, FUGW, or LentiCRISPRv2 (Addgene) (together with VSVG and psPAX2 plasmids). Viruses were harvested at 60-72 hours post transfection. And the viral supernatant was centrifuged at 600 g for 5 minutes to remove cell debris. The indicated cells were infected with the viral supernatant diluted with fresh medium (30% viral supernatant) with 10 µg/ml polybrene.

For the generation of TMED9-KO cell lines in Hela, the cells were infected with LentiCRISPRv2 virus containing TMED9 targeting sequences (sgRNA sequences: GTGCTGTGGCTGGCGACGCG & AGCGCGCTCTACTTTCACAT). Single colonies were isolated and determined for TMED9 KO.

### Cell-free LC3 lipidation, ERGIC isolation and immunoblot

These were performed as previously described^27,29^. Quantification of SEC12 relocation to the ERGIC was based on the percentage of ERGIC SEC12 relative to total SEC12. For immunoisolation of lipidated ERGIC, FLAG-tagged LC3 was used for the cell-free lipidation assay with the ERGIC membrane as shown before. Then agarose with anti-FLAG antibodies was employed to immunoisolate the lipidated ERGIC followed by FLAG peptide elution. The eluents was centrifuged at 100,000 xg to collect the isolated ERGIC membrane. Mass spectrometry was performed by the Protein Chemistry and Proteomics Center at Tsinghua University.

### Co-immunoprecipitation (Co-IP) and peptide pull-down assay

The details of co-IP was described before ^60,79^. Briefly, the indicated cells were lysed on ice for 30 min in IP buffer (50 mM Tris/HCl pH 7.4, 150 mM NaCl, 1 mM EDTA, 0.5% NP40, 10% glycerol) with protease inhibitors, and the lysates were cleared by centrifugation. The resulting supernatants were incubated with indicated agaroses and rotated at 4°C for 3 h. Then the agaroses were washed five times with IP buffer followed by immunoblot.

For peptide pull-down assay, 250 ug synthetic peptides (from Beijing Scilight Biotechnology LLC) were conjugated to 100 ul agarose beads using the AminoLink Plus Coupling Resin (Thermo, 20501) according to the manufacturers’ protocol. 2 μg purified FLAG-tagged SEC12 proteins were incubated with 15 μl peptides-coupled beads in Co-IP buffer and rotated at 4°C for 3 h. The agarose was then washed three times with Co-IP buffer. After washing, 2×SDS loading buffer was added to the beads, and immunoblot was performed as described previously^60,79^.

### Immunofluorescence microscopy and quantification

Immunofluorescence was described previously ^60,79^. In brief, the cells were incubated with 4% paraformaldehyde for 15 min at room temperature or cold methanol for 10 min (for SEC12 staining). The cells were permeabilized with 0.1% Triton X-100 or 50 μg/ml digitonin (for LC3 staining) diluted in PBS at room temperature for 3 min followed by incubating with 10% FBS diluted with PBS for 1 h and primary antibody incubation for 1 h. Cells were washed three times with PBS, followed by secondary antibody incubation for 1 h at room temperature. Fluorescence images were acquired using the Olympus FV3000 confocal microscope. Quantification was performed with using ImageJ as described previously ^60,79^.

### Determination of autophagic flux

For the evaluation of autophagic flux, Hela cells were infected with lentivirus containing mCherry-pHluorin-LC3B. After being treated with indicated conditions, the cells were fixed with 4% paraformaldehyde for 15 min at room temperature. Fluorescence images were acquired using the Olympus FV3000 confocal microscope. Autophagosome and autolysosome numbers were counted.

### Cycloheximide Chase experiments

For CHX chase assay, cells were transfected with SOD1 (G93A)-GFP plasmid. At 24 h after transfection, cells were treated with 50 μg/ml CHX, with or without 0.5 μg/ml Bafilomycin A1 as indicated and were collected at each indicated time point for immunoblot analysis.

### In vitro GUV contact assay

POPC (1-palmitoyl-2-oleoyl-glycero-3-phosphocholine), DOPE (1,2-dioleoyl-sn-glycero-3-phosphoethanolamine), POPS (1-palmitoyl-2-oleoyl-sn-glycero-3-phospho-L-serine), DOPE-rhodamine, PE-biotin and cholesterol were purchased from Avanti Polar Lipids. Lipids were mixed as POPC:DOPE:POPS:cholesterol (4:2.5:2.5:1). Using a glass syringe (HEMILTON) to add the lipid solution to the glass slide and allow the sample to dry with nitrogen gas for 3 h. 200 μl PBS was gently added to glass for 10 min to suspend the GUVs.

For the membrane tether assay, 2% PE-MPB (1,2-dioleoyl-sn-glycero-3-phosphoethanolamine-N-[4-(p-maleimidophenyl) butyramide]) was added to the “ERGIC-GUVs” lipid mixture. And 2% DGS-NTA (1,2-dioleoyl-sn-glycero-3-[(N-(5-amino-1-carboxypentyl) iminodiacetic acid) succinyl]) with 0.1% DOPE-rhodamine and 1% PE-biotin was added to the “ERES-GUVs” lipid mixture. FITC-TM9CT or FITC-TM9CT (m4) peptide was crosslinked to the “ERGIC-GUVs” and His-SEC12 was attached to the “ERES-GUVs” via His-Nickel interaction (300 μl solution contain 3 μg peptides or proteins). The crosslink or attachment reaction was performed for 30min with rotation at room temperature. In order to remove the free peptides or proteins, a membrane flotation procedure was performed. For each 300 μl solution, 300 μl 50% OptiPrep (diluted in PBS) was added. The mixture was overlaid with 480 μl 20% OptiPrep and 90 μl PBS, centrifuged at 100,000 xg for 2 h and the 150 μl top fraction (which contains the proteoliposomes) was collected. The “ERES-GUVs” and “ERGIC-GUVs” were mixed at a volume ratio of 1:1, and incubated for 30 min at room temperature. The mixtures were subsequently added to the streptavidin-coated glass cell culture dish and incubated for 20 min. The image of the GUVs was captured using a laser scanning confocal microscope (Olympus FV3000). Quantification was performed using Image J.

### In vitro transactivation SAR1 assay

For protein purification, genes encoding SAR1-BFP, SAR1-T39N-BFP and BFP were inserted into the PGEX4T1 vector. The proteins were expressed in *E. coli* BL21 at 22°C for 5 h. After expression, the bacteria were collected and digested with 0.5 mg/ml lysozyme (Sigma) in lysis buffer (50 mM Tris/HCl pH 8.0, 5 mM EDTA,150 mM NaCl,10% glycerol for GST protein purification) plus 0.3 mM DTT and protease inhibitors on ice for 0.5 h. Triton X-100 was added to adjust to 0.5% final concentration. The lysates were sonicated and centrifuged at 20,000 xg for 1 h. The supernatants were incubated with glutathione agarose rotated at 4°C for 2 h. The agarose was washed with 10 bed volume of wash buffer with 0.1% Tween20 and wash buffer each (PBS for GST protein purification). The proteins were eluted by elution buffers (50 mM Tris 8.0, 250 mM KCl, 25 mM glutathione for GST proteins). The proteins were snap frozen by liquid nitrogen and stored in PBS at 80°C. The purification of His-tagged human cytoplasmic domain SEC12 was performed as described previously^89^.

For the first assay described in the manuscript, we prepared small unilamellar vesicles (SUVs) as described previously^66^. The lipids mixture is the same as “ERGIC-GUVs” described above. The lipid mixtures were dried with a nitrogen stream and further dried for 1 h at 37°C. The lipid film was then suspended completely with PBS and subjected to 10 cycles of freezing in liquid nitrogen and thawing in a 42°C water bath. Finally, the liposomes were extruded 21 times through a 100 nm pore size polycarbonate film to produce the SUVs. The FITC-TM9CT peptides were added into the SUVs solution (300 μl solution contain 3 μg peptides) and incubated for 30 min with rotation at room temperature to crosslink. In order to remove the free peptides, a membrane flotation procedure was performed as described above. And the ERGIC-SUVs were the incubated with SEC12 crosslinked AminoLink-beads overnight at 4°C. The beads with the associated SUVs (beads-SUVs) were washed three times with PBS. After washing, 50 μg/ml SAR1-BFP or SAR1-T39N-BFP protein and indicated GTP were added to the beads-SUVs, and incubated at room temperature for 30 min. The beads-SUVs were washed three times with PBS. The ERGIC-SUVs bound to the SEC12 beads were eluted by incubating with 0.5 mg/ml of the TMED9-CT peptides for 30 min at room temperature followed by 1,500 xg centrifugation to separate ERGIC-SUVs and SEC12 beads. SDS loading buffer was added to the SEC12 beads and ERGIC-SUVs, and immunoblot was performed.

In the second assay described in the manuscript, generation of ERGIC-mimetic beads was based on a previous study^66^. In brief, The ERGIC-SUVs with 1% PE-biotin was incubated with Streptavidin agarose beads (GE) overnight at 4°C with slow rotation. The beads were washed three times with PBS. To test SAR1 recruitment to the ERGIC-mimetic beads, reaction mixes containing 0.15 mM GTP and SEC12 beads (described above) in a final volume of 200 µl were prepared. The reaction mixes were transferred to a confocal dish, and respective proteins (according to the experimental setup) were added at a final concentration of 50 μg/ml. Images were acquired after 1 h incubation 30°C in the dark using Olympus FV3000 confocal microscope with 60x objective and processed with Image J software.

### STORM and SIM analysis

Three-color STORM was performed on a homebuilt setup based on a Nikon Eclipse Ti-U inverted fluorescence microscope, as previously described^60^. Briefly, goat anti-mouse-Alexa Fluor 647 (Invitrogen A-21235), custom-labeled goat anti-rat-CF680, and custom-labeled goat anti-rabbit-CF568 were used as secondary antibodies for TMED9-V5, SEC12, and ERGIC-53, respectively. Alexa Fluor 647 and CF680 were simultaneously excited under 647-nm illumination and distinguished through ratiometric single-molecule detection using a dichroic mirror (Chroma T685lpxr), and CF568 was subsequently imaged with 560-nm excitation. For two color STORM, Dye-labeled cell samples were mounted on glass slides with a standard STORM imaging buffer consisting of 5% (w/v) glucose, 100 mM cysteamine, 0.8 mg/ml glucose oxidase, and 40 μg/ml catalase in Tris-HCl (pH 7.5)^90,91^. Imaging experiments were performed using a Nikon combined Confocal A1/SIM/STORM system with four activation/imaging lasers (405 nm, 488 nm and 561 nm from Coherent, 647 nm from MPBC) and a CFI Apo SR TIRF 100X oil (NA 1.49) objective. The images were acquired with an Andor EMCCD camera iXON 897. Data analysis was performed using the NIS-Elements AR (Nikon) software.

### Live cell imaging

Hela mCherry-SEC12 & EGFP-ERGIC53 cells were plated onto glass bottom cell culture dish (NEST, 801002). Before imaging, the medium was replaced with 2 ml of cell medium or starvation medium added as indicated. Individual culture dish was fitted into a heated stage on the microscope, and cells were maintained at 37°C. Images were acquired with the SD-SIM microscopy. The SD-SIM is a commercial system based on an inverted fluorescence microscope 461 (IX81, Olympus) equipped with a wide-field objective (100×/1.3 oil, Olympus) and a scanning confocal system (CSU-X1, Yokogawa). Four laser beams of 405 nm, 488 nm, 561 nm, and 647 nm were combined with the SD-SIM. The Live-SR module (GATACA systems, France) was equipped with the SD-SIM. The images were captured by an sCMOS camera (C14440-20UP, Hamamatsu, Japan). A 3D surface model was generated, and quantification of the ERES-ERGIC contact was carried out using Imaris 9 software.

### Electron microscopy and electron tomography

Hela cells were fixed with 2.5% glutaraldehyde for 1h at room temperature and washed 3 ×15 min with 0.1 M PB. Post-fixation staining was performed with 1% osmium tetroxide (SPI, 1250423) for 30 min on ice. Cells were washed 3 ×15 min with ultrapure water, and then placed in 1% aqueous uranyl acetate (EMS, 22400) at 4°C overnight. Samples were then washed 3 ×15 min with ultrapure water, and dehydrated in a cold-graded ethanol series (50%, 70%, 80%, 90%, 100%, 100%, 100%; 2min in each). Penetrating in EPON 812 resin using 1:1 (v/v) resin and ethanol for 8 h, 2:1 (v/v) resin and ethanol for 8 h, 3:1 (v/v) resin and ethanol for 8 h, then pure resin 2 × 8 h and finally into fresh resin and polymerization in oven at 60°C for 48 h. Embedded samples were sliced into 75-nm-thick sections and stained with uranyl acetate and lead citrate (C1813156). Samples were imaged under the HT-7800 120kv transmission electron microscope.

For electron tomography, samples for TEM were prepared as described above. The samples of ROI were cut into 250-nm-thick sections. The grids were imaged on a F20 electron microscope (Thermo Fisher Scientific, Hillsboro, OR) operated at a voltage of 200 kV and K2 direct electron detector (Gatan, CA). Cellular organelles of interest were recorded in counting mode at a nominal magnification of 32,000 x, resulting in a calibrated pixel size of 1.533 nm. Tilt-series were collected using continuous scheme from -61° to 60° at 1° steps and defocus around -2 um. Tile-series are aligned by using Etomo patch tracking and relative tomograms are reconstructed by weighted back projection with SIRT-like filter. The ERES, ERGIC and proximal vesicles were modeled in IMOD. The heat map representing the distance between ERES and ERGIC were calculated based on the 3D modeling of two organelles.

For DAB staining and preparation of cultured cells for EM, transfected U2OS cells were fixed using room temperature 2% glutaraldehyde in buffer (100 mM sodium cacodylate with 2 mM CaCl_2_, pH 7.4), then quickly moved to ice. Cells were kept between 0 and 4°C for all subsequent steps until resin infiltration. After 30–60 min, cells were rinsed 5 × 2 min in chilled buffer, treated for 5 min in buffer containing 20 mM glycine to quench unreacted glutaraldehyde followed by 5 × 2 min rinses in chilled buffer. A freshly diluted solution of 0.5 mg/ml (1.4 mM) DAB tetrahydrochloride or the DAB free base (Sigma) dissolved in HCl was combined with 0.03% (v/v) (10 mM) H_2_O_2_ in chilled buffer, and the solution was added to cells for 5 min. The generation of reaction product could be monitored by transmitted LM. To halt the reaction, the DAB solution was removed, and cells were rinsed 5 × 5 min with chilled buffer. Post-fixation staining was performed with 2% osmium tetroxide for 30 min in chilled buffer. Cells were rinsed 5 × 2 min in chilled distilled water and placed in chilled 2% aqueous uranyl acetate overnight. The samples were then dehydrated in a cold graded ethanol series (20%, 50%, 70%, 90%, 100%, 100%) 2 min each, rinsed once in room temperature anhydrous ethanol to avoid condensation, and infiltrated in Durcupan ACM resin (Electron Microscopy Sciences) using 1:1 (v/v) anhydrous ethanol and resin for 30 min, and 100% resin 2 × 1 h. Finally the samples were embeded into fresh resin and polymerized in a vacuum oven at 60 °C for 48 h. DAB-stained areas of embedded cultured cells were identified by transmitted light, and the areas of interest were sawed out using a jeweler’s saw and mounted on dummy acrylic blocks with cyanoacrylic adhesive (Krazy Glue, Elmer’s Products). The coverslip was carefully removed, the block trimmed, and ultrathin (80 nm thick) sections were cut using an ultramicrotome (Leica Ultracut UTC6). Samples were imaged under the HT-7800 120kv transmission electron microscope.

### Secretion analysis

For determination of cargo secretion, cells were transfected with ssGFP. At 24 h after transfection, HEK293T cells were replaced with DMEM for 1 h. The medium was concentrated (20-fold) by a 10 kD Amicon filter (Millipore) and cell lysate was collected. Immunoblot was performed to determine the amount of cargoes in the medium and cell.

### RUSH system

Hela cells and HEK293T cells were cultured and infected with the RUSH reporter SBP-HA-SEC12 by lentivirus. To release the RUSH reporters from the ER, 40 µM biotin (Sigma, B4501) was added to the cultured cells at the indicated time. Images (Hela) were acquired by a Nikon combined Confocal A1/SIM/STORM system. Membrane fractionation and immunoblot (HEK293T) was performed as described above^27,29^.

Table S1 The lists of proteins identified in mass spectrometry as shown in **Figure 1C**

The proteins identified were listed in the table including names, gene ID, number of peptides, location and annotation of cytoplasmic length.

Table S2 Information about the antibodies used in this work and list of the siRNAs used in this study

Video S1 SD-SIM analysis of HeLa cells incubated in nutrient-rich medium (NR) related to **Figure 3D**

Video S2 SD-SIM analysis of HeLa cells starved (ST) in EBSS related to **Figure 3D**

Video S3 SD-SIM analysis of HeLa cells transfected with control siRNAs and starved (ST) in EBSS related to **Figure 3F**

Video S4 SD-SIM analysis of HeLa transfected with siRNAs against TMED9 and starved (ST) in EBSS related to **Figure 3F**

Video S5 Tomogram related to **Figure 3H**

---

**Figure S1.**
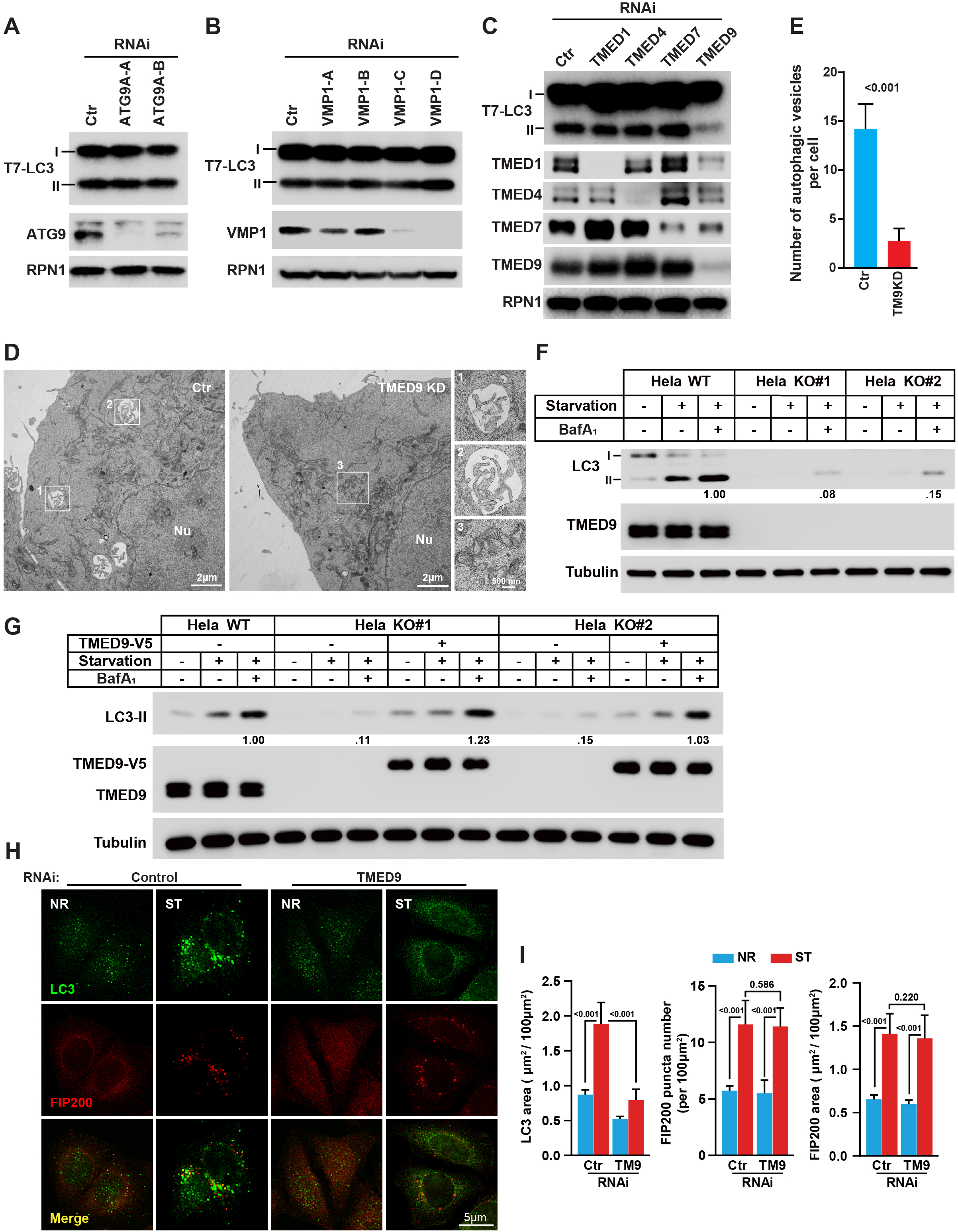
TMED9 regulates autophagosome biogenesis. (A-C) The membrane fraction from Atg5KO MEF cells transfected with control and siRNAs against ATG9A (A), VMP1 (B), or TMED1, TMED4, TMED7, TMED9 (C) was collected. Cell-free lipidation was performed with the Atg5KO MEF membranes and cytosols prepared from starved HEK293T cells. The blots are representative of at least three independent experiments (D) EM showing the autophagic vesicles in HeLa cells (control or TMED9 KD) starved in EBSS for 1 h. Scale bar sizes are indicated in the picture. (E) Quantification of the number of autophagic vesicles per cell analyzed in (D). Error bars represent standard deviations of >30 cells from three experiments. P-values obtained from two-tailed t-test. (F) LC3 lipidation in HeLa cells (control or TMED9 knockout (KO)). The cells were incubated in nutrient-rich medium or starved in EBSS for 1 h in the absence or presence of 500 nM bafilomycin A1 for 1 h. Immunoblots were performed to determine the levels of indicated proteins. Quantification was performed similarly to Fig.1G. The blots are representative of at least three independent experiments. (G) LC3 lipidation in HeLa cells (control or TMED9 KO) with or without TMED9-V5 re-expression. The cells were incubated in nutrient-rich medium or starved in EBSS for 1 h in the absence or presence of 500 nM Bafilomycin A1 for 1 h. Immunoblots were performed to determine the levels of indicated proteins. Quantification was performed similarly to Fig.1G. The blots are representative of at least three independent experiments. (H) Immunofluorescence of HeLa cells (control, TMED9 KD) with V5 and FIP200 antibodies. The cells were incubated in nutrient-rich medium or starved in EBSS for 1 h. (I) Quantification of the LC3 puncta area (μm^2^ / 100 μm^2^ cell area, left), FIP200 puncta number per 100μm^2^ cell area (middle) and FIP200 puncta area (μm^2^ /100 μm^2^ cell area, right) analyzed in (H). Error bars represent standard deviations of >150 cells from three independent experiments (>50 cell per experiment). P-value was obtained from two-tailed t-test.

**Figure S2.**
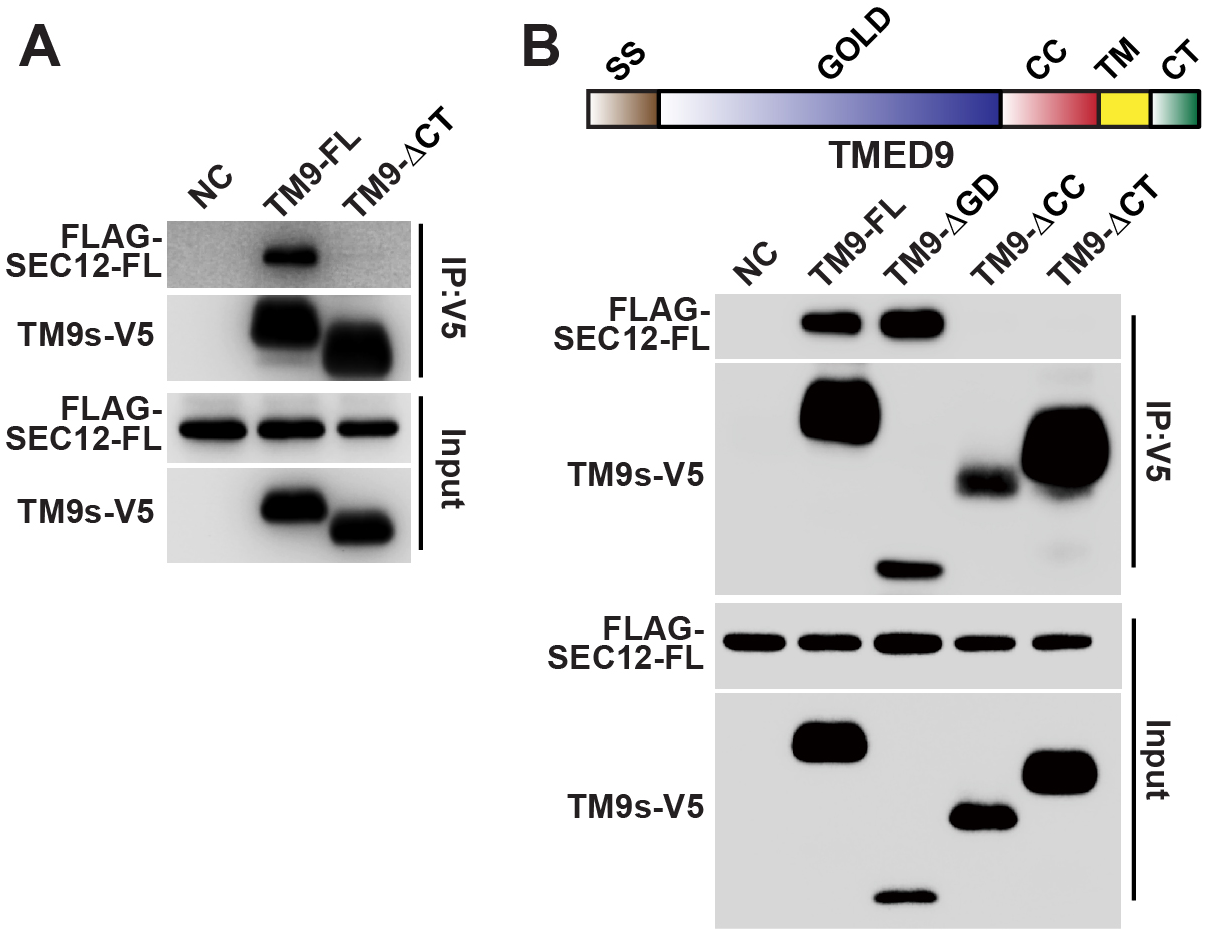
TMED9 interacting with C-terminal of SEC12 via the C-terminal cytoplasmic tail. (A) CoIP analysis of TMED9-V5 variants (FL and ΔCT) with FLAG-SEC12 in HEK293T starved in EBSS for 1 h using anti-V5 agarose. The blots are representative of at least three independent experiments. (B) CoIP analysis of TMED9-V5 variants (FL, ΔGOLD, ΔCC, and ΔCT) with FLAG-SEC12 in HEK293T starved in EBSS for 1 h using anti-V5 agarose. The blots are representative of at least three independent experiments.

**Figure S3.**
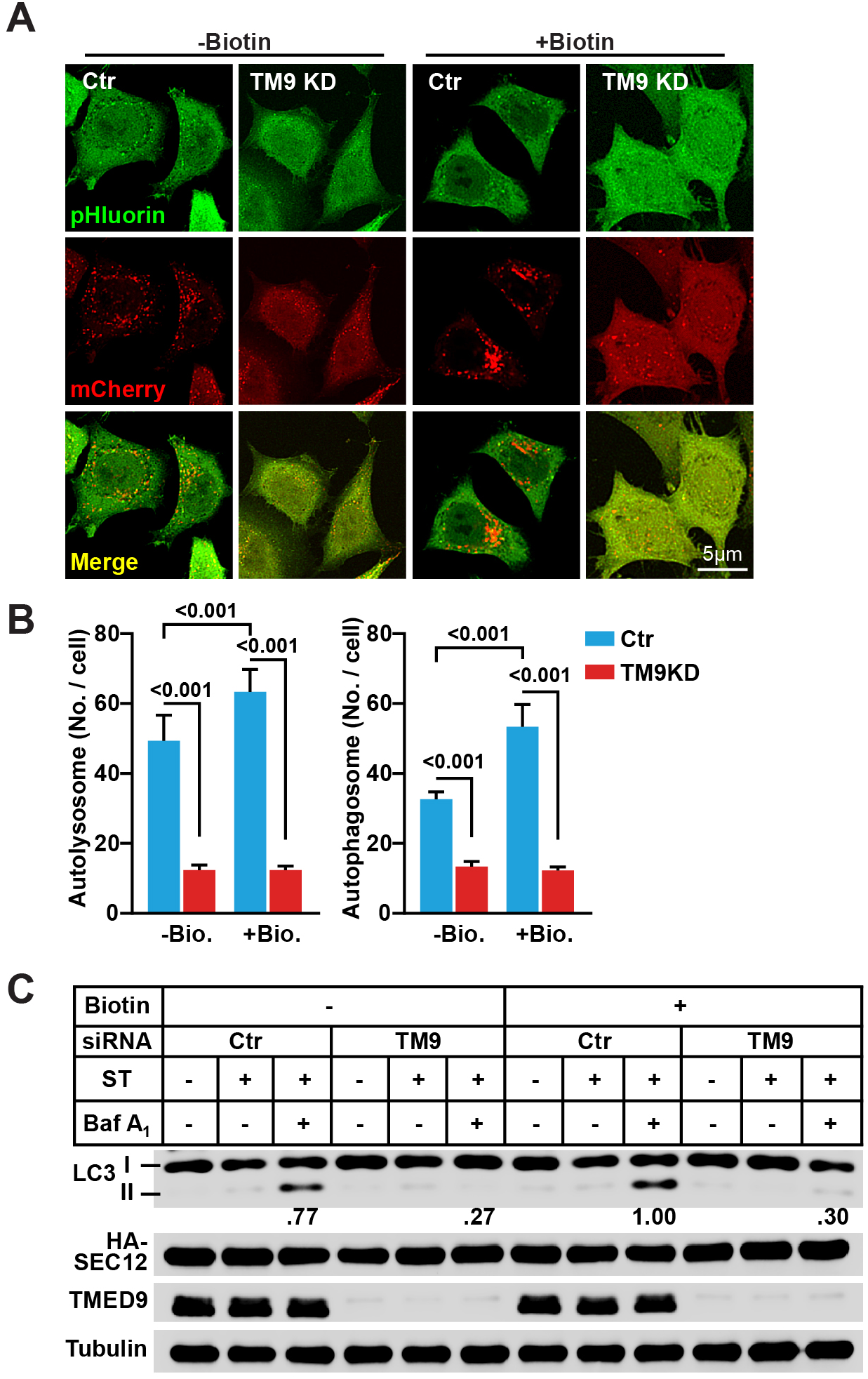
Inhibiting SEC12 relocation to the ERGIC marginally affects autophagosome biogenesis. (A) HeLa cells stably expressing the mCherry-pHluorin-LC3B (control and TMED9 KD) were treated with or without 40 μM biotin for 1 h and starved in EBSS for 1 h. Confocal microscopy was performed. (B) Quantification of autolysosome (red, left) and autophagosome (yellow, right) numbers per cell as shown in (A). Error bars represent standard deviations of >300 cells from three independent experiments (>100 cell per experiment). P-value was obtained from two-tailed t-test. (C) LC3 lipidation in HeLa cells stably expressing RUSH-SEC12 (SEC12 KD) transfected with control or siRNAs against TMED9. The cells were treated with or without 40 μM biotin for 1 h and incubated in nutrient-rich medium or starved in EBSS in the absence or presence of 500 nM bafilomycin A1 for 1 h. Immunoblots were performed to determine the levels of indicated proteins. Quantification was performed similarly to Fig.1G. The blots are representative of at least three independent experiments.

